# Dietary Oxysterols Reprogram Hepatic Lipid Metabolism and Reshape the Gut Metabolome–Microbiome Interface

**DOI:** 10.64898/2026.04.12.717948

**Authors:** Lisaura Maldonado-Pereira, Thuraya M. Mutawi, Arunima Singh, Brian J. Sanderson, Michaella J. Rekowski, Carlo Barnaba, Ilce G. Medina Meza

## Abstract

Dietary oxysterols are biologically active cholesterol oxidation products ubiquitous in Western diets, yet their systemic effects on host metabolism and the gut microbiome remain largely unexplored. Here, we employed an integrated multi-omics approach – shotgun metagenomics, quantitative proteomics, untargeted metabolomics, and bulk RNA-seq – to characterize the impact of DOxS exposure on the gut–liver axis in rats fed a Western diet (WD vs. WD-DOxS). Hepatic proteomics revealed near-complete suppression of the mevalonate/cholesterol biosynthesis pathway, particularly in males, while de novo lipogenesis enzymes (Scd1, Fasn, Plin2) were paradoxically upregulated, consistent with dual oxysterol signaling through SREBP inhibition and LXR activation. Bile acid synthesis was concurrently suppressed, confirmed by metabolomics. Strikingly, RNA-seq across liver, heart, and brain detected virtually no differentially expressed genes, establishing that DOxS act predominantly through post-transcriptional mechanisms. In the gut, DOxS increased microbial α-diversity while depleting *Limosilactobacillus reuteri*, with concomitant loss of the barrier-protective metabolite 3-indoleacrylic acid. Tissue-specific responses were widespread, with liver and colon frequently mounting opposing metabolic and immune responses to the same dietary challenge. Cross-omics integration revealed convergent microbiome–metabolite axes connecting microbial remodeling to both hepatic lipid reprogramming and colonic barrier disruption. These findings reposition dietary oxysterols from food-quality markers to active modulators of the gut–liver axis, with implications for metabolic disease and intestinal barrier integrity.

## INTRODUCTION

The Western Diet (WD) is characterized by an abundance of “*ready-to-eat”, “ready-to-drink”, “fast”,* and *“convenient”* products grouped as ultraprocessed foods (UPFs) according to NOVA.^1^ Up to 73% of the foods within the WD are UPFs.^2^ The production of UPFs alters the food’s structure, nutritional content, and taste,^3–5^ and growing epidemiological evidence links high UPF consumption to the onset of chronic disorders, including cardiometabolic disease, cancer, and neurodegeneration.^6^ Several mechanisms explain the link between increased consumption of UPFs and the development of chronic disorders. First, UPFs often replace minimally processed foods, reducing the intake of beneficial micronutrients, antioxidants, and phytochemicals.^7^ Nutrients in UPFs are largely acellular, which enhances nutrient availability in the small intestine, thereby promoting inflammation and gut dysbiosis.^8, 9^ Second, UPFs contribute to the overconsumption of total calories.^10^ Third, they expose individuals to non-nutritive food substances that have been associated with gut inflammation in preclinical studies.^6^ Fourth, a growing body of epidemiological evidence supports the hypothesis that UPFs consumption is detrimental to the gut microbiome.^11, 12^ Thus, understanding how UPFs contribute to chronic disorders through these mechanisms is crucial for developing effective dietary interventions and improving public health outcomes.

Among the chemical species generated during food processing, dietary oxysterols ^13^ are particularly relevant. DOxS are chemical species unavoidably generated during industrial food manufacturing, including light exposure, packaging, and storage.^13–15^ DOxS preserve the steroidal motif of cholesterol with an additional hydroxyl, ketone, or epoxy group.^16^ Over 100 DOxS are identified, including 7α/β-hydroxysterols^17^, 7-ketocholesterol (7-keto), 5,6α/β-epoxysterols,^13, 15^ cholestane-triol (triol), and 20-hydroxycholesterol (20-OH) are the most abundant in foods. Our group has established DOxS as quantitative biomarkers of UPF quality, building the FooDOxS database spanning over 1,680 food items across 25 countries.^13^ *In vitro*, DOxS exert well-documented pro-inflammatory, pro-oxidant, and pro-apoptotic effects.^18–20^ At the molecular level, oxysterols are potent regulators of cholesterol homeostasis through their interactions with SREBP–Insig–SCAP complexes and liver X receptors (LXRs),^21–23^ placing them at the intersection of lipid metabolism, inflammatory signaling, and bile acid synthesis. Despite this mechanistic potency, the systemic consequences of chronic dietary DOxS exposure in vivo – particularly their impact on the gut microbiome and the gut–liver metabolic axis – have not been characterized.

Standard high-fat diet ^24^ models do not capture the complexity of the Western diet, as they lack refined sugars, processed ingredients, and food additives that characterize human UPF consumption. The Total Western diet (WD), developed by Hintze and co-workers^25, 26^ to better model human dietary patterns, uses a combination of high-fat, high-sugar, low-fiber, and industrially processed ingredients (such as anhydrous milk fat) to mimic the metabolic and inflammatory effects observed in humans consuming typical Western diets. However, neither HFD nor WD includes DOxS in their formulation criteria, leaving a critical dietary component unaccounted for. Herein, we compared the WD with a modified WD formulation supplemented with DOxS (WD-DOxS) at exposure levels informed by our FooDOxS database.^13^ Using a rat model with a 2×2 factorial design (diet × sex), we first confirmed that DOxS are absorbed and detected in plasma with distinct pharmacokinetic profiles. We then conducted a multi-week dietary intervention and applied an integrated multi-omics framework – shotgun metagenomics, quantitative proteomics, untargeted metabolomics, and bulk RNA-seq – to characterize the systemic impact of DOxS across the gut–liver axis. Our results reveal that dietary DOxS profoundly restructures the cecal microbiome, increasing community diversity while depleting key barrier-protective commensals and their tryptophan-derived metabolites. In the liver, DOxS triggered a coordinated suppression of cholesterol biosynthesis alongside an activation of de novo lipogenesis, operating predominantly through post-transcriptional mechanisms that rendered the hepatic transcriptome largely unresponsive. Tissue-specific and often opposing responses in liver and colon underscore the complexity of DOxS-mediated metabolic reprogramming. Collectively, these findings reposition dietary oxysterols from markers of food processing to biologically active modulators of the gut–liver axis, with implications for understanding how ultra-processed Western diets contribute to metabolic disease.^6^

## MATERIALS AND METHODS

### Experiment 1. Kinetics of absorption and excretion of DOxS in rats

Animal protocol was approved by the Animal Committee (MSU PROTO202100168) at Michigan State University. Male Wistar rats (6-8 weeks) n=8 per group were purchased from Charles River Laboratories®. After acclimatization (1 week) with AIN93G diet, rats were randomly divided into 3 groups and administered by oral gavage individual DOxS using 0.3 mL in olive oil as vehicle. A side chain oxysterol (25-hydroxycholesterol) and a ring B oxysterol (7β-hydroxycholesterol) were used in addition to cholesterol-D-7. Blood was collected by venipuncture at baseline, 1, 2, 3, 4, 6, 8, 12, and 24 h. At the end of the experiments, rats were euthanized, and blood at the terminal point was collected for further analysis.

### Experiment 2. Effect of DOxS chronic exposure on gut microbiota

Male (n=10) and female (n=10) Wistar rats (6-8 weeks) were purchased from Charles River Laboratories®. After 1 week of acclimatization, rats were divided into two groups (G1: Western Diet, G2: WD-DOxS) with 5 females and 5 males per group. Diets were administered for 6 weeks *ad libitum*. Weight was monitored; blood and stools were collected weekly. At the end of the experiment, rats were euthanized, and the liver, colonic, and cecal tissues were harvested and snap frozen for further analysis. An animal from G2 (WD-DOxS) group did not survive the full dietary study and was not included for further analysis.

### Diets design and formulation

Diets were custom-formulated by Inotiv (Envigo Teklad, Madison, WI), on an energy density basis, as described elsewhere.^25, 26^ This approach allows for modeling human nutrient intakes on a caloric basis by taking into account the different energy needs of rodents vs. humans. <u>Western Diet (WD):</u> TB 230378 Description. Designed based on customer specifications for nutrient values based on Pre-Pandemic NHANES mean data adjusted on a caloric basis. Modification of TD.110424 with ∼1.4% increase in fat by weight. The fatty acid profile is approximately 34.6% SUFA, 40.7% MUFA, and 24.7% PUFA. Choline increased by 2% by weight. Pink color added for visual distinction. Both experimental diets were formulated to replicate median daily nutrient intake levels for US individuals >2 years based on NHANES 2017-2018 survey.^27^

#### WD-DOxS Diet

This diet maintains identical macronutrient composition to the WD diet but supplemented with a mixture of DOxS standards (≥99% purity; Avanti Polar Lipids), at physiologically relevant concentrations based on our FooDOxS database and exposure assessment models for UPFs intake.^13, 14^ The DOxS mixture comprises the five most abundant in UPFs: 7-ketocholesterol (60.9 μg/g), 7β-hydroxycholesterol (60.9 μg/g), 5,6α-epoxycholesterol (40.6 μg/g), Cholestanetriol (20.3 μg/g) and 20-hydroxycholesterol (20.3 μg/g), ratios prevalence in the >80% of UPFs. Diets were obtained in a pellet, not irradiated by the vendor, stored under gaseous nitrogen, and maintained at 4°C to avoid further oxidation. After formulation but before use in this study, the DOxS profile for all experimental diets was verified according to Maldonado et al.^28^ Diets were color-coded (pink for WD, orange for WD + DOxS) to facilitate experimental blinding and accurate feeding protocols.

### Shotgun metagenomics

Cecal content was flushed using sterile PBS, collected, weighed, and snap-frozen before being shipped to CosmosID for next-generation sequencing analysis. DNA from cecal samples was isolated using the QIAGEN DNeasy PowerSoil Pro Kit, according to the manufacturer’s protocol. DNA samples were quantified using Qubit 4 fluorometer and Qubit™ dsDNA HS Assay Kit (Thermofisher Scientific). DNA libraries were prepared using the Watchmaker DNA Library Prep Kit. Genomic DNA was fragmented using a mastermix of Watchmaker Frag/AT Buffer and Frag/AT Enzyme Mix. IDT xGen UDI Primers and IDT Stubby Adapters were added to each sample followed by 7 cycles of PCR to construct the DNA libraries. The final DNA libraries were purified using AMPure magnetic beads (Beckman Coulter) and eluted in nuclease-free water. Following elution, the libraries were quantified using the Qubit™ fluorometer dsDNA HS Assay Kit. Libraries were then sequenced on the Element AVITI platform 2×150 bp.

#### Bioinformatics Analysis via CosmosID-HUB Methods

The patented Kepler multi-kingdom taxonomic profiler is a comprehensive algorithm consisting of three interwoven pipelines: (1) pre-computation database construction, (2) K-mer based taxonomic identification and classification, and (3) abundance estimation and classification refinement.

1. Taxonomic classification relies on the CosmosID GenBook database, a curated collection of ∼30,000 species spanning Bacteria, Archaea, Fungi, Protists, and Viruses. Genomes are selected based on assembly completeness, low contamination, and intra-species diversity, then filtered to remove low-complexity sequences, prophages, plasmids, and host-derived regions. Curated genomes are decomposed into variable-length n-mers and organized into a phylogenetic tree structure, where shared biomarkers form the backbone and species-unique biomarkers occupy the leaves.
2. K-mer based taxonomic classification and identification. This first comparator phase splits the millions of sequencing reads for each sample into k-mer sets. It then looks for exact matches between sample k-mers and reference biomarkers that are dispersed across the different phylogenetic branches and leaves of the GenBook™ database. By focusing on small genomic regions of large differential informational content, this phase eliminates 99% of genomes that are almost certainly not in the metagenomic sample and generates a short list of nearest reference strains. This highly efficient and accurate classification readily admits sub-species taxonomy to an ANI divergence less than 0.003. Classification sensitivity and accuracy are maintained through biomarker aggregation statistics, coverage depth estimation, and abundance estimation-but with an inhomogeneous variance that is parsed into Step (3).
3. Probabilistic Smith-Waterman-based abundance Estimation and classification refinement. The second comparator phase uses an edit distance-scoring based probabilistic Smith-Waterman algorithm to compare sequencing reads with a reference set of identified microbial taxa from Step (2). This involves a more computationally-intensive processing of the remaining 1% of taxa to eliminate any false positives and achieve abundance estimations that are more homogeneous, more accurate, and have significantly less variance. Take an example scenario of 10 reads, where reads 1-8 belong to taxon A, read 9 belongs to taxon B, and read 10 belongs to both taxon A and taxon B with equal alignment score. Kepler will probabilistically split the abundance of read 10 into taxon A with an abundance of 8.89 and taxon B with an abundance of 1.11. Given a more complicated metagenomic sample scenario, contested reads are apportioned according to implied priors from uncontested reads. An iterative Maximum Likelihood Estimation algorithm is then converged to the final read abundance estimates. To summarize, overall abundance precision and classification accuracy achieved by running the comparators in sequence, scoring the entire read probabilistically against the reference set, and distinguishing homologous regions.

#### Functional Analysis

Initial QC, adapter trimming and preprocessing of metagenomic sequencing reads are done using BBduk.^29^ The quality-controlled reads are then subjected to a translated search against a comprehensive and non-redundant protein sequence database, UniRef 90. The UniRef90 database, provided by UniProt,^30^ represents a clustering of all non-redundant protein sequences in UniProt, such that each sequence in a cluster aligns with 90% identity and 80% coverage of the longest sequence in the cluster. The mapping of metagenomic reads to gene sequences are weighted by mapping quality, coverage and gene sequence length to estimate community wide weighted gene family abundances as described by Franzosa et al.^31^ Gene families are then annotated to MetaCyc (4) reactions (Metabolic Enzymes) to reconstruct and quantify MetaCyc^32^ metabolic pathways in the community as described by Franzosa et al.^31^ Furthermore, the UniRef_90 gene families are also regrouped to Enzyme Commission Enzymes, Pfam protein domains, CAZy enzymes and GO Terms to get an exhaustive overview of gene functions in the community. Lastly, to facilitate comparisons across multiple samples with varying sequencing depths, the abundance values are normalized using Total Sum Scaling (TSS) normalization to produce “Copies per million” (analogous to transcripts per million, or TPMs, in RNA-Seq) units.

To independently validate CosmosID findings, raw reads were reprocessed using two complementary profiling strategies. Marker-gene-based taxonomy was performed with MetaPhlAn 4,^33^ and k-mer-based taxonomy with Kraken 2^34^ followed by Bayesian abundance re-estimation with Bracken.^35^ Quality control and adapter trimming were performed with fastp,^36^ and host (rat) reads were removed by mapping against the mRatBN7.2 reference genome^37^ using Bowtie2.^38^ Differential abundance was assessed using MaAsLin2^39^ with diet as the primary variable and sex as a covariate. Results were independently validated using LEfSe^40^ with default parameters (alpha = 0.05, LDA threshold = 2.0). Alpha diversity (Chao1, Shannon, Simpson) was compared between groups using the Mann-Whitney U test, and beta diversity (Bray-Curtis, abundance-weighted Jaccard) was evaluated by PERMANOVA with 999 permutations. P-values were corrected for multiple testing using the Benjamini-Hochberg method (q < 0.05).

### Tissues transcriptomics

A total of 57 RNA samples were obtained from rat brain, liver, and heart tissues. Animals were divided into two groups: a control group maintained on a Western diet (n = 10; 5 males and 5 females) and an experimental group receiving a Western diet supplemented with DOxS (n = 9; 5 males and 4 females). Total RNA was extracted using the Zymo Research RNA extraction kit with on-column DNase I treatment. RNA concentration and integrity were assessed with the Qubit RNA High Sensitivity Assay Kit (Thermo Fisher Scientific) and Agilent TapeStation, respectively. mRNA libraries were prepared from all 57 samples using the NEBNext Ultra II Directional RNA Library Prep Kit for Illumina (New England Biolabs) with poly(A) selection. Indexed libraries were evaluated on the TapeStation, pooled equimolarly to 2 nM (average insert size: 289 bp), and purified by bead-based size selection. Sequencing was performed on an Illumina NextSeq 2000 platform (NX2K-P4-SR100, X-LEAP chemistry) using single-end 100 bp reads, yielding an average of ∼18 million reads per sample.

### Differential Expression Analysis

Raw sequencing reads were processed using the University of Kansas Genomic Data Science Core RNA-seq pipeline v. 0.1.0 using Nextflow v. 24.10.3.^41, 42^ Reads were trimmed for adapter sequences and for minimum quality scores using fastp v. 0.23.4 with the parameters -q 15 and -u 40 (**Table S1**). Trimmed reads were then aligned to the rat genome assembly GRCr8 (GCA_036323735.1) with the Ensembl version 114 annotation using STAR v. 2.7.9a, and read count quantification was performed by RSEM v. 1.3.1 (**Table S2**).^43^ The resulting counts matrix was then used to estimate patterns of differential gene expression using R v.4.4.1.^44^ The count data were first inspected for batch effects using plotMDS v. 3.60.5 (**Fig. S1**).^45^ Patterns of differential gene expression were modeled for each tissue separately in DESeq2 v. 1.44.0 with two sets of models: 1) using the formula counts ∼ treatment + sex + treatment × sex (Table S3), and 2) using the formula counts ∼ treatment + sex (no interaction effect; Table S4). The resulting log_2_ fold-change values were adjusted using the DESeq2 function lfcShrink with the ashr method.^46^ Genes were classified as differentially expressed based on thresholds of Benjamini-Hochberg FDR-adjusted *P-*values (*P_FDR_*) < 0.01 and absolute log_2_ fold-change values > 2. Plots were generated with ggplot2 v. 3.5.1,^47^ cowplot v. 1.1.3.

### Mass spectrometry proteomics

Colon and Liver tissue samples were homogenized by transferring them into homogenization tubes along with 30 μL of RIPA buffer with protease and phosphatase inhibitors and nuclease per mg of tissue and using the Beat homogenizer on standard power for two 10-minute cycles. Samples were centrifuged at 14000 g for 10 mins at 4°C and homogenates were transferred to new tubes. BCA assay was performed, and based on the concentrations, 20 µL of protein from each sample was used for the remainder of the sample preparation. Samples volumes were increased to 50 uL using 50 mM TEAB. Disulfides were reduced by adding 5 µL of 50 mM TCEP (5mM final) and incubating at 55 °C for 30 minutes. Alkylated cysteines by adding 1.5ul of 375mM IAA (10mM final) and incubating at RT in the dark for 30 minutes. Precipitated proteins by adding 225 uL of acetone (1:5 dilution) and incubating at −20C overnight. Centrifuged samples at 14000 g for 10 minutes at 4°C. Pipetted off supernatant and allowed proteins to air dry on the bench top for 15 minutes. Proteins were resuspended in 100 uL of 50mM TEAB, 2mM CaCl_2_, and digested by adding 500 ng of trypsin (0.1 mg/mL) and incubating overnight at 37 °C at 500 RPM. Reactions were quenched by the addition of 10% formic acid to 1%. Peptide concentrations were measured using the nanodrop UV system, and based on the concentrations 1-3 μL of each sample was loaded onto the C18 RP column and injected into the Orbitrap Ascend mass spectrometer with FAIMS for LC-MS/MS analysis. Data was searched using the Proteome Discoverer SeQuest software against the rat database downloaded from UniProt on 9-12-2025.

### Proteomics Data Processing and Differential Expression Analysis

Over-Representation Analysis (ORA) was performed separately on upregulated and downregulated DEP lists for each comparison using clusterProfiler v4.12^48^ in R v4.3. Enrichment was tested against Gene Ontology Biological Process (GO-BP), Gene Ontology Molecular Function (GO-MF), KEGG,^49^ and Reactome^50^ databases using the rat annotation package. The full set of quantified proteins for each comparison was used as the background universe. *P*-values were adjusted for multiple testing using the Benjamini-Hochberg procedure; terms with adjusted *p* < 0.05 were considered significant. Gene Set Enrichment Analysis (GSEA) was performed on the full ranked protein list for each comparison using clusterProfiler.^48^ Proteins were ranked by the metric sign (log₂FC) × −log₁₀(*p*-value), which integrates both the direction and statistical significance of expression changes. Enrichment was tested against GO-BP, KEGG, and Reactome gene sets with minimum and maximum gene set sizes of 15 and 500, respectively. An epsilon value of 0 was used to enable exact *p*-value computation. The 50 Hallmark gene sets from the Molecular Signatures Database^51^ (MSigDB) were obtained for *Rattus norvegicus* via ortholog mapping using the msigdbr package v10.0.^52^ Gene set enrichment was performed using fgsea v1.30^53^ with 10,000 permutations. Normalized enrichment scores (NES) were computed for each comparison, and results were considered significant at adjusted *p* < 0.05. A cross-comparison NES heatmap was generated to visualize pathway-level responses across all four tissue × sex combinations. Transcription factor (TF) activities were inferred from the log₂FC proteomics data using decoupleR v2.10^54^ with the univariate linear model method. F-target regulons were obtained from DoRothEA,^55, 56^ filtered to confidence levels A, B, and C (representing interactions supported by curated literature, ChIP-seq, and/or transcription factor binding motif evidence), comprising 271 TFs and 13,223 interactions. Because the proteomics data are from rat, gene symbols were converted to uppercase human orthologs before regulon matching, which is standard practice for DoRothEA-based analyses. TFs with fewer than 5 targets in the dataset were excluded. Activity scores were tested for significance and corrected using the Benjamini-Hochberg method. Volcano plots were generated using matplotlib v3.9^57^ with proteins color-coded by functional category: cholesterol biosynthesis, de novo lipogenesis/lipid metabolism/bile acid metabolism, xenobiotic/cytochrome P450 metabolism, immune/interferon signaling, and antioxidant response. Pathway enrichment heatmaps were generated using pheatmap^58^ in R.

### Metabolomics

Previously homogenized colon and liver tissue from proteomics analysis were extracted by diluting the homogenate 4:1 in a solution of 40% methanol, 40% acetonitrile, 20% 50 mM ammonium bicarbonate. Samples were incubated for 30 min on ice to precipitate proteins and then centrifuged at 14,000 x *g* for 10 min at 4°C. The supernatant containing the polar metabolite fraction was transferred to a new vial and dried in a vacuum centrifuge. Samples were stored at −20C until analysis. Samples were resuspended in 0.1% formic acid and injected in duplicate on the tims TOF HT (Bruker®) equipped with an Elute HPLC (Bruker). Briefly, the extracted metabolite sample was injected onto an Intensity Solo C18-2 (2.1 x 100 mm, Bruker) at a flow rate of 0.25 mL/min at 1%B (0.1% FA in acetonitrile, mobile phase A: 0.1% FA in water) and held for 2 min prior eluting with a linear gradient from 1-99% B in 17 min. The column was washed for 2 min at 99% B before re-equilibration in 1% B for 2 min to prepare for the next injection. The column was heated to 35°C. Metabolites were ionized in positive (4500 V) or negative (−4500 V) mode using the same MS method. Briefly, data were collected between *m/z* 80-1300 in PASEF mode (1) with 1/K_0_ values 0.45-1.45 V*s/cm^2^ with a ramp time of 100 ms. Two PASEF ramps were collected, and metabolites were fragmented by CID at 50% NCE. Chemical class enrichment was assessed by assigning metabolites to chemical classes based on structural annotation and testing for directional class-level shifts using the Wilcoxon signed-rank test on log₂FC values within each class. All statistical analyses were performed in Python 3.11 using SciPy v1.13 and statsmodels v0.14.

### Statistical Analysis

Two-sided Mann-Whitney U tests were used for all pairwise comparisons unless otherwise specified. Multiple testing correction employed the Benjamini-Hochberg (BH) FDR procedure; significance was defined as padj < 0.05 with |fold change| > 1.5 (|log2FC| > 0.585) for metabolomics and metagenomics, and p < 0.05 with |log2FC| > 0.585 for proteomics DEPs (uncorrected, as ratios and p-values were computed within Proteome Discoverer). Outliers exceeding 2 SD from the group mean were excluded from pharmacokinetic analyses. Beta diversity significance was assessed by PERMANOVA and ANOSIM (999 permutations). MaAsLin2 was the primary model for metagenomics species-level associations, with sex included as a covariate; LEfSe served as validation. Pathway and gene set enrichment analyses used clusterProfiler^48^ with BH correction, and MSigDB Hallmark sets were tested via fgsea.^53^ Transcription factor activity was inferred using decoupleR^54^ with the ULM method and DoRothEA regulons (confidence levels A–C).^55, 56^ Sex was treated as a biological variable; all analyses were stratified by sex rather than pooled. Statistical analyses were performed in Python 3.11 (SciPy, statsmodels) and R 4.3 (clusterProfiler, fgsea, decoupleR). All reported p-values are two-sided.

## RESULTS

### Experiment 1. DOxS have different absorption profiles from cholesterol

To confirm systemic bioavailability, a preliminary pharmacokinetic study was conducted in male Wistar rats (n = 8) receiving individual DOxS by oral gavage in olive oil. Plasma concentrations of cholesterol-*d7*, 25-hydroxycholesterol (25-OH), and 7β-hydroxycholesterol (7β-OH) were monitored over 24 h (**Fig. 1A**). Plasma cholesterol declined progressively from 4.29 ± 0.29 µg/mL at baseline to 2.68 ± 0.19 µg/mL at 24 h (−37.7%; P < 0.001), with levels significantly below baseline from 3 h onward (P = 0.042) (**Fig. 1B**). 25-OH was undetectable at baseline, appeared in circulation within 1 h (0.13 ± 0.01 µg/mL; P = 0.002), peaked at 3 h (0.63 ± 0.15 µg/mL), and remained significantly elevated at 24 h (0.29 ± 0.10 µg/mL; P < 0.001), indicating rapid absorption and sustained systemic exposure (**Fig. 1C**). 7β-OH rose more gradually from 0.16 ± 0.02 µg/mL at baseline to a peak of 0.37 ± 0.02 µg/mL at 6 h and remained elevated at 24 h (0.27 ± 0.04 µg/mL; P = 0.018) (**Fig. 1D**). These results confirmed that individually administered DOxS are rapidly absorbed and sustain measurable plasma levels over 24 h, supporting the rationale for chronic dietary exposure.

**Figure 1.**
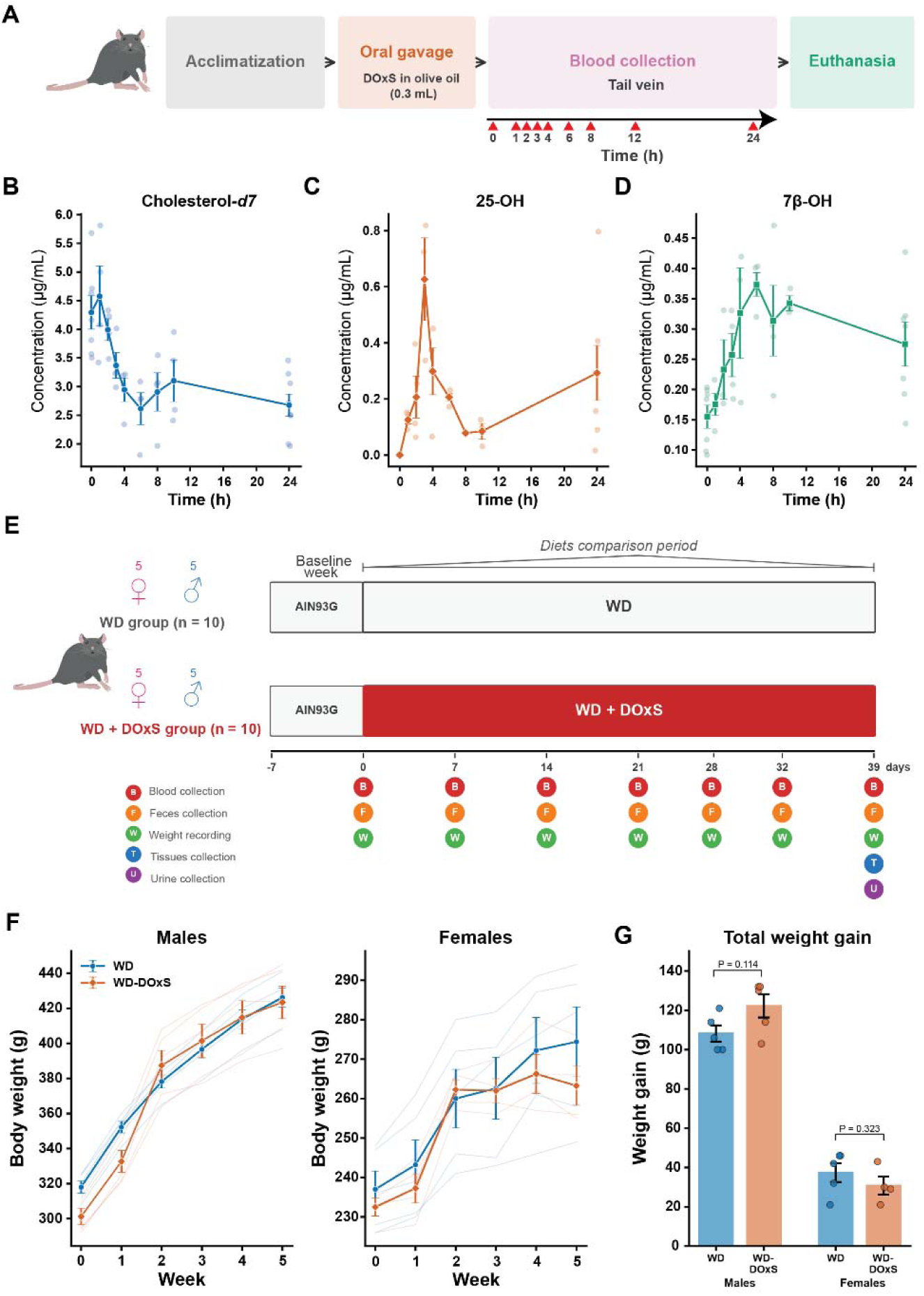
DOxS are systemically bioavailable and do not alter body weight in rats fed a Western diet. (**A**) **Exp. 1.** Schematic of the pharmacokinetic study design. Male Wistar rats (n = 8) received individual DOxS by oral gavage; blood was collected from the tail vein at baseline and 1-24 h post-administration. **(B–D)** Plasma concentrations of cholesterol-*d7* (B), 25-hydroxycholesterol (25-OH; C), and 7β-hydroxycholesterol (7β-OH; D) over 24 h. Lines connect group means ± SEM; individual data points are shown. **(E) Exp. 2.** Schematic of the dietary intervention study. Male and female Wistar rats were randomized to a Western diet (WD; n = 10) or WD supplemented with DOxS (WD + DOxS; n = 10) following a one-week acclimatization and one-week baseline period on AIN93G diet. **(F)** Body weight trajectories for males and females (right). Thin lines represent individual animals; thick lines show group means ± SEM. **(G)** Total weight gain (final − baseline) by diet and sex. Bars represent mean ± SEM with individual data points. All P-values (Mann-Whitney U test) are two-sided.

### Experiment 2. Body weight

Having established systemic bioavailability, we next examined the effects of chronic DOxS exposure by feeding rats a Western diet supplemented with DOxS (WD-DOxS) or a Western diet alone (WD) for 6 weeks. Male rats gained substantially more weight than females over the study period, consistent with normal sexual dimorphism (**Fig. 1F**). Within each sex, DOxS exposure did not significantly alter body weight trajectories or total weight gain (**Fig. 1F-G**). Males in the WD-DOxS group gained 122.2 ± 5.9 g compared with 108.2 ± 4.1 g in WD (Mann-Whitney U, P = 0.114), while females gained 30.8 ± 4.6 g versus 37.4 ± 4.8 g, respectively (P = 0.323) (**Fig. 1G**). These results indicate that DOxS exposure at the administered dose did not affect growth.

### Diet drives distinct community composition

Shotgun metagenomic sequencing of cecal contents yielded a total of 463.2 million paired-end reads across 19 samples (n = 10 WD; n = 9 WD-DOxS). Mean sequencing depth was 24.8 ± 4.6 million reads per sample in the WD group and 30.0 ± 10.4 million in the WD-DOxS group, with no significant difference between groups (Mann-Whitney U test, P = 0.167), indicating that sequencing effort did not confound downstream comparisons. Taxonomic and functional profiling was performed using the CosmosID platform. To assess whether DOxS exposure alters within-sample microbial diversity, we computed three complementary alpha diversity metrics. All three were significantly elevated in WD-DOxS relative to WD (**Fig. 2A**). Chao1 richness, an estimate of total species count, was 101.6 ± 22.8 in WD-DOxS compared with 64.2 ± 10.6 in WD (P = 0.005). The Simpson index, which emphasizes dominance, was 0.86 ± 0.05 versus 0.77 ± 0.08. Shannon diversity, which accounts for both richness and evenness, was 3.78 ± 0.65 versus 2.74 ± 0.41 (P = 0.021). Together, these results indicate that WD-DOxS supports a richer and more even microbial community than WD alone. When stratified by sex, no significant sex effect was observed on any alpha diversity metric (all P > 0.67, Mann-Whitney U test), confirming diet as the primary driver of within-sample diversity (**Fig. S1A**). We next evaluated whether the two diets harbored compositionally distinct microbial communities using beta diversity analysis. Principal coordinates analysis (PCoA) of both abundance-weighted Jaccard and Bray-Curtis distance matrices revealed clear separation between WD and WD-DOxS samples (**Fig. 2B**). PERMANOVA confirmed that diet explained a significant fraction of community variation for both metrics (Jaccard: F = 4.60, P = 0.001; Bray-Curtis: F = 5.87, P = 0.001; 999 permutations). These results demonstrate that DOxS exposure reshapes the overall structure of the gut microbiome.

**Figure 2.**
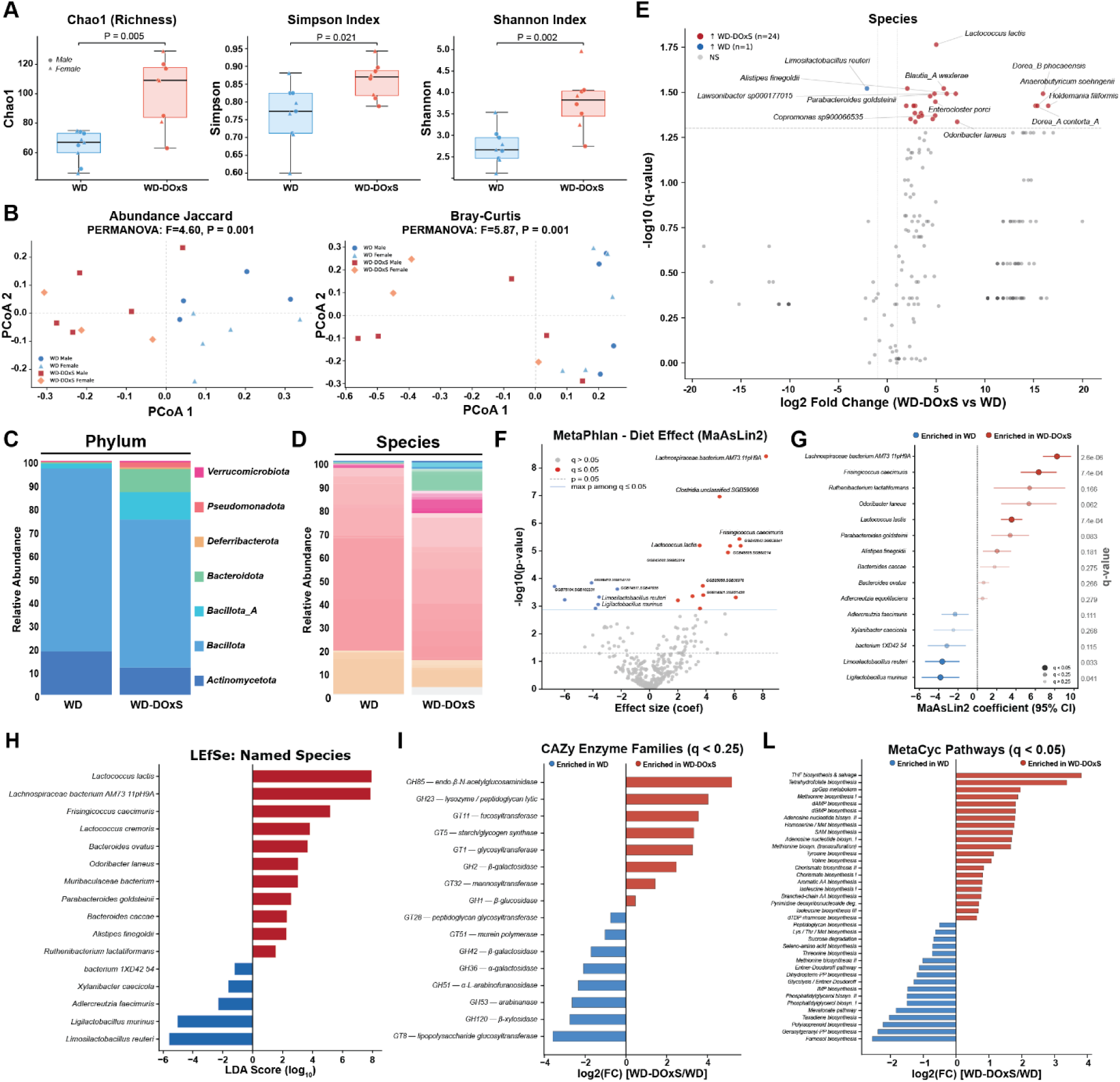
DOxS exposure reshapes gut microbiome composition and function, promoting taxonomic expansion and divergent metabolic pathway enrichment. **(A)** Alpha diversity metrics. Boxplots show median and interquartile range; individual animals are plotted by sex (circle = male, triangle = female). P-values from Mann-Whitney U tests. **(B)** Principal coordinates analysis (PCoA) of abundance-weighted Jaccard and Bray-Curtis (right) distance matrices, colored by diet and sex. PERMANOVA statistics are shown above each plot. **(C)** Phylum-level relative abundance averaged by diet group. **(D)** Species-level relative abundance stacked bar plots by diet group. **(E)** Volcano plot of differentially abundant species (Mann-Whitney U, Benjamini-Hochberg correction). **(F)** MaAsLin2 analysis of diet effect on species abundance controlling for sex, showing effect size versus −log10(P-value). Dashed lines indicate significance thresholds. **(G)** Forest plot of MaAsLin2 coefficients (95% CI) for species significantly associated with diet. **(H)** LEfSe analysis of named species, ranked by LDA score. Bars indicate the diet group in which each taxon is enriched. **(I)** Differentially abundant CAZy enzyme families (q < 0.25), shown as log2 fold change (WD-DOxS vs. WD). **(L)** Differentially abundant MetaCyc pathways (q < 0.05), shown as log2 fold change. All P-values are two-sided; multiple comparisons were corrected using Benjamini-Hochberg FDR.

### WD-DOxS promotes expansion of Bacillota and Bacteroidota taxa

At the phylum level, both diets were dominated by *Bacillota* ^59^ and *Bacteroidota*, with visible shifts in their relative proportions between groups (**Fig. 2C**). Differential abundance testing at the species level (Mann-Whitney U test with Benjamini-Hochberg correction) identified 25 species as significantly different between diets (q < 0.05) (**Fig. 2D**). Strikingly, 24 of 25 were enriched in WD-DOxS, while only a single species-*Limosilactobacillus reuteri-* was enriched in WD (log_2_FC = −2.09, q = 0.030; **Fig. 2E**). Among WD-DOxS-enriched species, the largest effect sizes were observed for *Holdemania filiformis* (log_2_FC = +16.5), *Dorea phocaeensis* (+16.0), and *Anaerobutyricum soehngenii* (+15.4), which were essentially absent from WD samples. More moderately enriched taxa included *Alistipes finegoldii* (+7.0), *Parabacteroides goldsteinii* (+6.1), *Blautia wexlerae* (+5.8), and *Ruminococcus gnavus* (+4.6). Taxonomically, the enriched species spanned *Bacillota/Bacillota_A* (15 of 24), *Bacteroidota* (7 of 24), and *Actinomycetota* (2 of 24), indicating that WD-DOxS promotes a broad taxonomic expansion rather than a phylum-specific bloom. Notably, several WD-DOxS-enriched taxa are recognized producers of short-chain fatty acids ^60^ or their metabolic precursors. *Anaerobutyricum soehngenii* is a well-characterized butyrate and propionate producer that uniquely converts lactate and acetate to butyrate.^61, 62^ *Blautia wexlerae* produces succinate, lactate, and acetate, and its oral administration has been shown to ameliorate obesity and type 2 diabetes in mice via metabolic remodeling of the gut microbiota.^63^ *Alistipes finegoldii* is an emerging strain known to produce succinic acid as its major fermentation end product, with minor amounts of acetic and propionic acid.^64^ *Parabacteroides goldsteinii* primarily generates acetate and succinate and has demonstrated anti-obesity effects in high-fat diet–fed mice through SCFA-mediated pathways and enhanced intestinal integrity.^65^ The sole WD-enriched species, *Limosilactobacillus reuteri*, is a well-characterized lactate producer commonly associated with high-sugar, low-fiber dietary conditions.^66^

To independently validate these findings, we conduct a secondary analysis of all samples using an open-source pipeline employing two complementary classification approaches: marker-gene-based (MetaPhlAn 4)^33^ and k-mer-based (Kraken2/Bracken) profiling,^67, 68^ with differential abundance assessed using MaAsLin2^39^ with sex included as a covariate (**Fig. 2F-G**). This analysis identified 20 species significantly associated with diet (q < 0.05) out of 337 tested, with 13 enriched in WD-DOxS and 7 in WD. No species showed a significant sex effect after multiple testing correction. The strongest WD-DOxS associations included *Lachnospiraceae bacterium* AM73 11pH9A (coef = 8.23, q < 0.001), *Frisingicoccus caecimuris* (coef = 6.35, q < 0.001), and *Lactococcus lactis* (coef = 3.55, q < 0.001). Consistent with CosmosID profiling, *L. reuteri* was again depleted in WD-DOxS (coef = −3.63, q = 0.033), alongside a second *Lactobacillaceae* member, *Ligilactobacillus murinus* (coef = −3.81, q = 0.041), which was identified only after controlling for sex. Similarly, *Parabacteroides goldsteinii* (q = 0.083) and *Odoribacter laneus* (q = 0.062) showed consistent DOxS enrichment across platforms, though at relaxed significance thresholds. Several unclassified species-level genome bins (SGBs) within the *Clostridia* and *Lachnospiraceae* were among the most strongly affected taxa, suggesting that DOxS exposure also shapes microbial lineages not yet represented in current reference databases. These results were further corroborated by LEfSe analysis^40^ (**Fig. 2H**), which identified 76 differentially abundant species (48 enriched in WD-DOxS, 28 in WD) and ranked *Lactococcus lactis* (LDA = 7.94) and *Lachnospiraceae* bacterium AM73 11pH9A (LDA = 7.87) as the strongest discriminators for WD-DOxS, with *L. reuteri* (LDA = 5.57) and *L. murinus* (LDA = 5.01) as the top WD-associated species, fully consistent with the MaAsLin2 results. Collectively, DOxS exposure was associated with depletion of *Lactobacillaceae* and expansion of *Lachnospiraceae*, *Bacteroidota*, and SCFA-producing taxa; these patterns were consistently observed across all three profiling platforms and confirmed using two independent statistical frameworks.

### Gene-level functional enrichment favors WD despite lower diversity

To determine whether the taxonomic shifts translated to functional consequences, we performed differential abundance testing across six annotation layers: CAZy carbohydrate-active enzyme families^69^ (**Fig. 2I**), EC enzyme numbers, GO molecular functions, GO biological processes, Pfam protein families (**Fig. S2B-D**), and MetaCyc metabolic pathways^32^ (**Fig. 2L**). In contrast to the species-level pattern, the functional layers revealed a more balanced picture. Across all six layers, 866 features were differentially abundant (q < 0.05), of which 476 (55.0%) were enriched in WD versus 390 (45.0%) in WD-DOxS (CAZy 8, EC 118, GO-MF 186, GO-BP 58, Pfam 458, MetaCyc 38; **Supplementary Fig. S2B-D**). This apparent decoupling between taxonomic diversity and functional enrichment suggests that the WD microbiome, while harboring fewer species, carries a more concentrated functional repertoire per taxon. Conversely, the greater taxonomic diversity under WD-DOxS distributes gene content across a broader set of organisms, potentially diluting per-taxon functional signal when measured as relative abundance. Notably, GO molecular function analysis revealed that antioxidant activity was significantly enriched in the WD-DOxS microbiome (log2FC = +1.38, q = 0.015), while GO biological process analysis identified glutathione biosynthesis as enriched in WD (log2FC = −1.87, q = 0.021), suggesting that the two communities employ distinct redox defense strategies in response to dietary oxidized cholesterol.

### WD-DOxS activates amino acid and folate biosynthesis pathways

To move beyond individual genes toward interpretable metabolic context, we analyzed MetaCyc pathway-level abundances. Of 342 detected pathways, 38 were differentially abundant between diets (q < 0.05): 21 enriched in WD-DOxS and 17 in WD (**Fig. 2I**). Pathways enriched in WD-DOxS were dominated by amino acid biosynthesis, including branched-chain amino acid pathways (valine biosynthesis, log2FC = +1.08; isoleucine biosynthesis I and III, +0.77 and +0.68; branched-chain AA biosynthesis, +0.76), aromatic amino acid pathways (chorismate biosynthesis, +0.84; aromatic AA biosynthesis, +0.80; tyrosine biosynthesis, +1.14), and methionine/SAM-cycle pathways (methionine biosynthesis I, +1.88; SAM biosynthesis, +1.72; homoserine/Met biosynthesis, +1.77). Tetrahydrofolate biosynthesis pathways showed the largest effect sizes of any functional category (THF biosynthesis, log2FC = +3.37; THF biosynthesis & salvage, +3.81). Nucleotide biosynthesis (dAMP and dGMP biosynthesis, both +1.81) and ppGpp metabolism (+1.96), a stringent response regulator associated with bacterial growth-state transitions,^70^ were also elevated. Pathways enriched in WD were characterized by isoprenoid and terpenoid biosynthesis, including farnesol biosynthesis (−2.56), geranylgeranyl-PP biosynthesis (−2.38), polyisoprenoid biosynthesis (−2.23), and the mevalonate pathway (−1.83). Phosphatidylglycerol biosynthesis (both isoforms, −1.50), the Entner-Doudoroff pathway (−1.12), and sucrose degradation (−0.68, q = 0.007) were also enriched. This functional signature points to a WD community investing in membrane lipid remodeling^71^ and alternative carbon catabolism.^72^ Together, the pathway-level analysis reveals a functional divergence that complements the taxonomic shifts: WD-DOxS drives a biosynthetically active community oriented toward amino acid production, one-carbon metabolism, and nucleotide synthesis, whereas the WD community emphasizes isoprenoid-based membrane maintenance and alternative glycolysis.

### RNAseq Differential Expression

To evaluate whether DOxS exposure induces transcriptional changes across host tissues, we performed bulk RNA-seq on liver, heart, and brain from all animals in both diet groups. Cecal tissue was excluded from transcriptomic analysis due to insufficient RNA integrity, a common challenge in luminal tissues with high endogenous RNase activity.^73^ Our RNA-seq libraries generated an average of 32,450,006 ± 5,606,406 reads per sample, of which an average of 75.59% mapped uniquely to the reference transcriptome (**Table S1**). Initial assessment of raw gene expression counts with multidimensional scaling (MDS) plots showed that samples clustered well by tissue (**Fig S2A**). Because most of the variation in the full data set is between tissues, we assessed differential expression separately for each tissue. Assessments of raw gene expression counts with MDS plots within tissues showed that samples clustered by sex and treatment (**Fig S2B**), although one sample from brain tissue was an outlier relative to the others and was thus excluded from downstream analyses (sample Gp1_WD_brain, **Fig S2C**). Across all tissues, we found very few genes that exhibited significant differences in expression due to the main effect of treatment with DOxS, either as a main effect or a sex-by-treatment interaction effect.

Due to the very small number of significant genes with a sex-by-treatment interaction, as well as visual inspection of the results, we used separate models to estimate the genes with significant interactions (**Table S3, Supplemental Material**) and then used models without the interaction term to estimate the genes with significant main effects of DOxS treatment (**Table S4, Supplemental Material**) within each tissue. In the liver, treatment with DOxS led to a significant increase in expression of *Rfx6* in females, and a significant decrease in *Sephs2* expression in females (**Fig S2D**; **Table S3, Supplemental Material**). Additionally, there was an overall decrease in expression in *Sycp3l2*, although this pattern appears to be entirely driven by four female libraries (despite the sex-by-treatment interaction not being significant; **Fig S2D**; **Tables S3-4, Supplemental Material**). In the brain, there were no genes with a significant sex-by-treatment interaction, but *Mug1* and *Apoa4* showed overall decreases in expression in response to DOxS treatment, while *Rps2-ps43* showed increased expression (**Fig S2E**; **Tables S3-4, Supplemental Material**). In the heart, DOxS treatment drove increased expression of *Moxd1*, *Cr2*, and *Cd19*, though this pattern was only evident in a small number of samples (**Fig S2F**; **Tables S3-4, Supplemental Material**).

### Quantitative Proteomics Reveals Tissue- and Sex-Specific Metabolic Reprogramming by DOxS exposure

To investigate the molecular mechanisms through which DOxS alter host tissue biology in the context of a Western Diet, we performed quantitative shotgun proteomics on colon and liver tissues. Colon and liver were selected as target tissues given their respective roles as the primary site of dietary oxysterol exposure along the gut–liver axis and the central organ of systemic cholesterol metabolism,^74^ and their known susceptibility to oxidative and lipotoxic stress.^15^ Protein identification and quantification were performed using Proteome Discoverer with Sequest HT, yielding 7,295 high-confidence proteins (FDR < 1%). Differentially expressed proteins (DEPs) were defined as those with a fold change ≥ 1.5 (|log2FC| > 0.585) and a p-value < 0.05. Each comparison assessed the effect of DOxS exposure within a given tissue and sex (i.e., WD-DOxS/WD). DOxS exposure induced widespread proteomic changes in both tissues, with a general bias toward downregulation (**Fig. 3**). In the colon, 1,375 DEPs were identified in females (434 upregulated, 941 downregulated) and 1,149 in males (320 upregulated, 829 downregulated) (**Fig. S3A**). The liver exhibited a more balanced profile, with 905 DEPs in females (402 upregulated, 503 downregulated) and 949 in males (426 upregulated, 523 downregulated) (**Fig. S3A**). Cross-sex comparison within each tissue revealed substantial sex-specific responses: in the colon, only 556 DEPs were shared between males and females, of which 401 were directionally concordant and 155 were discordant (i.e., upregulated in one sex and downregulated in the other) (**Fig. S3B**). In the liver, 328 DEPs were shared, with 196 concordant and 132 discordant (**Fig. S3B**). Cross-tissue overlap was even more limited, with only 290 shared DEPs in females and 224 in males, indicating that DOxS exerted predominantly tissue-specific effects on the proteome (**Fig. 3A**).

**Figure 3.**
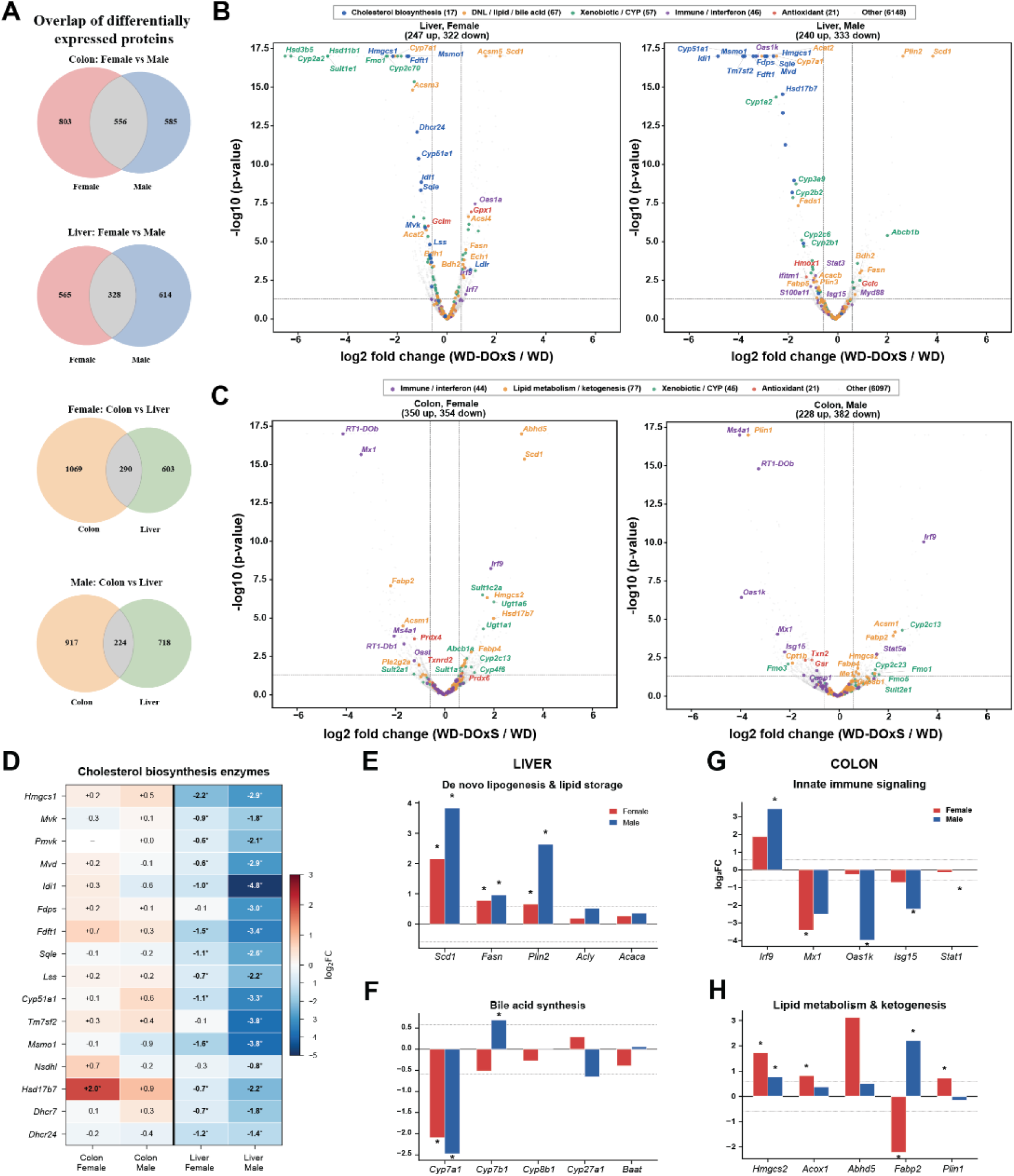
DOxS exposure suppresses hepatic cholesterol biosynthesis and activates tissue-specific lipid remodeling and immune signaling programs. **(A)** Venn diagrams showing overlap of differentially expressed proteins (DEPs; p < 0.05, |log2FC| > 0.58) between sexes within each tissue (top) and between tissues within each sex (bottom). **(B)** Volcano plots of liver DEPs in females and males (right). Proteins are colored by functional category. Total upregulated and downregulated counts are indicated. **(C)** Volcano plots of colon DEPs in females and males (right), colored by functional category. **(D)** Heatmap of log2 fold changes for cholesterol biosynthesis pathway enzymes across all four tissues × sex comparisons. Asterisks denote statistical significance (q < 0.05). **(E)** Log2 fold change of de novo lipogenesis and lipid storage proteins in liver, stratified by sex. **(F)** Log2 fold change of bile acid synthesis enzymes in liver. **(G)** Log2 fold change of innate immune signaling proteins in colon. **(H)** Log2 fold change of lipid metabolism and ketogenesis proteins in colon, stratified by sex. Bars represent mean log2FC (WD-DOxS vs. WD); asterisks indicate q < 0.05. DEPs were defined as proteins with |log2FC| > 0.585 and p < 0.05. DEPs were defined as |log2FC| > 0.585 and p < 0.05. Proteins identified at 1% FDR.

### DOxS cause suppression of the liver cholesterol biosynthesis pathway

The most striking finding of the hepatic proteome analysis was a near-complete downregulation of the mevalonate–cholesterol biosynthesis pathway in DOxS-exposed animals. This effect was observed in both sexes but was markedly more pronounced in males. Of 16 enzymes spanning the pathway from HMG-CoA synthase to the terminal reductases, 14 were significantly downregulated in males and 12 in females (**Fig. 3B**). The magnitude of suppression in males was significant: Idi1 (isopentenyl-diphosphate delta isomerase 1; log2FC = −4.84, p < 10–17), Msmo1 (methylsterol monooxygenase 1; −3.83), Tm7sf2 (Δ-14-sterol reductase; −3.76), Fdft1 (squalene synthase; −3.41), Cyp51a1 (lanosterol 14α-demethylase; −3.30), Fdps (farnesyl diphosphate synthase; −2.95), and Hmgcs1 (HMG-CoA synthase 1; −2.88) all exhibited reductions exceeding 4-to 29-fold. In females, the same enzymes were significantly downregulated but at lower magnitudes (log2FC range: −0.61 to −2.18). Importantly, none of these enzymes were significantly altered in the colon in either sex, establishing the liver-specific nature of this response (**Fig. 3C**, **Fig. S4A**). This pattern is consistent with a feedback inhibition model in which exogenous oxysterols suppress SREBP2 processing,^75^ thereby coordinately downregulating the transcription of cholesterol biosynthetic genes.^21^ In contrast to the suppression of cholesterol biosynthesis, DOxS exposure was associated with a significant upregulation of key de novo lipogenesis (DNL) enzymes in the liver. Stearoyl-CoA desaturase 1 (Scd1) was the most consistently and dramatically upregulated protein, with log2FC values of +3.83 in males (p < 10–17) and +2.14 in females (p < 10–17), corresponding to approximately 14-fold and 4-fold increases, respectively. Fatty acid synthase (Fasn) was also significantly upregulated in both sexes (log2FC = +0.96 in males, p = 7.5 × 10−4; +0.77 in females, p = 3.4 × 10−5). Additionally, the lipid droplet coat protein perilipin 2 ^76^ was markedly elevated, particularly in males (log2FC = +2.63, p < 10–17), suggesting increased hepatic lipid droplet accumulation. Scd1 was also upregulated in the colon of females (log2FC = +3.24, p < 10–15) but not males, pointing to a sex-specific cross-tissue effect. The concurrent upregulation of fatty acid synthesis with suppression of cholesterol synthesis suggests that DOxS may redirect hepatic lipid metabolism away from cholesterol toward triglyceride and fatty acid accumulation, a pattern with potential implications for hepatic steatosis. Cholesterol 7α-hydroxylase, the rate-limiting enzyme in the classical bile acid synthesis pathway,^23^ was significantly downregulated in the liver in both females (log2FC = −2.09, p < 10–17) and males (log2FC = −2.47, p < 10–17), representing approximately 4-to 5.5-fold reductions. This is mechanistically consistent with the known capacity of oxysterols to suppress Cyp7a1 through activation of LXR/FXR signaling axes.^23^ The suppression of bile acid synthesis, in conjunction with the mevalonate pathway downregulation, implies a broad inhibition of hepatic cholesterol utilization pathways by dietary DOxS. Beyond the cholesterol- and bile acid-metabolizing CYPs, DOxS exposure was associated with broad downregulation of drug- and xenobiotic-metabolizing cytochrome P450 enzymes in the liver (**Fig. S4B-D**). Among the most affected were Cyp1a2 (log2FC = −2.49 in males, −0.67 in females), Cyp2b2 (−1.81 in males, −1.32 in females), Cyp2c6 (−1.46 in males), Cyp2c11 (−1.84 in females), Cyp2c70 (−2.41 in females), and Cyp2a2 (−6.29 in females), all reaching statistical significance. The suppression exhibited notable sex specificity, with certain isoforms preferentially affected in one sex over the other (e.g., Cyp2c11 in females only, Cyp2c6 in males only). Of 72 CYP/SULT/UGT family members quantified, the predominant direction in the liver was downregulation. This pattern raises potential concerns regarding the capacity of DOxS-exposed animals to metabolize xenobiotics and pharmaceuticals, warranting further pharmacokinetic investigation.

The hepatic antioxidant response to DOxS exposure was mixed and sex-dependent. In females, glutathione peroxidase 1 (Gpx1; log2FC = +0.98, p = 1.2 × 10−7) and glutathione S-transferase Pi 1 (Gstp1; +0.64, p = 1.4 × 10−4) were significantly upregulated, suggesting an adaptive antioxidant response. However, glutathione S-transferase Mu 1 (Gstm1) was significantly downregulated in females (−0.93, p = 3.1 × 10−7). In males, NAD(P)H quinone dehydrogenase 1 (Nqo1; −0.71, p = 0.017) and heme oxygenase 1 (Hmox1; −1.27, p = 1.9 × 10−3) were significantly downregulated. The absence of a coordinated upregulation of the classical Nrf2-mediated antioxidant defense in males, despite the oxidative challenge posed by DOxS, may reflect an impaired adaptive capacity (**Fig. S4C**).

### DOxS exposure causes altered innate immune signaling and lipid metabolism in colon

The colon exhibited a distinctive pattern of innate immune modulation following DOxS exposure. Interferon regulatory factor 9 (Irf9) was strongly upregulated in both females (log2FC = +1.88, p = 5.9 × 10−9) and males (+3.44, p = 8.9 × 10–11). Paradoxically, several canonical interferon-stimulated genes ^77^ were downregulated, particularly in males: Mx1 (myxovirus resistance protein 1; −2.50, p = 9.2 × 10−5), Oas1k (2’-5’-oligoadenylate synthetase; −3.97, p = 3.7 × 10−7), and Isg15 (ubiquitin-like modifier; −2.21, p = 1.3 × 10−3). In females, Mx1 was similarly reduced (−3.40, p < 10–15) (**Fig. S3C**). This uncoupling of Irf9 upregulation from ISG expression suggests a potential disruption of the JAK-STAT–ISGF3 signaling axis at a step downstream of Irf9 ^78,78^ or the engagement of non-canonical Irf9 functions independent of the classical type I interferon response.^77, 79^ (**Fig. 3G**).

The colon displayed a pattern of lipid metabolic alterations distinct from the liver (**Fig. S5**). Mitochondrial HMG-CoA synthase (Hmgcs2), the rate-limiting enzyme for ketogenesis, was significantly upregulated in both females (log2FC = +1.72, p = 4.7 × 10−7) and males (+0.76, p = 0.015), while it was unchanged in the liver. Peroxisomal acyl-CoA oxidase 1 (Acox1), involved in fatty acid β-oxidation, was upregulated in female colon (+0.82, p = 8.5 × 10−3). The lipase co-activator Abhd5 was markedly upregulated in female colon (+3.12, p < 10–17) and modestly in liver females ^80^. A notable sex-discordant response was observed for intestinal fatty acid-binding protein (Fabp2), which was significantly downregulated in females (−2.20, p = 7.9 × 10−8) but upregulated in males (+2.20, p = 1.2 × 10−4) (**Fig. 3H; Fig. S3C**). Unlike the liver, the cholesterol biosynthesis pathway was not significantly altered in the colon, indicating that the oxysterol-mediated suppression of mevalonate enzymes was restricted to the liver.

### Sex Dimorphism in the Proteomic Response

Sex-dependent differences were widespread across the dataset and were evident at both the individual protein and pathway levels. Beyond the quantitative differences noted above (e.g., larger magnitudes of cholesterol pathway suppression in males), there were qualitative differences in which proteins and programs were affected. In the colon, 147 of 556 shared DEPs were discordant in direction between males and females; in the liver, 132 of 328 were discordant, representing discordance rates of 26% and 40%, respectively (**Fig. S5**). Hallmark GSEA revealed sex-divergent pathway activation in the colon: females showed significant enrichment of Estrogen Response Late (NES = +1.78, padj = 0.024) and a positive trend in IL2-STAT5 Signaling (NES = +1.67, padj = 0.084), while males showed suppression of E2F Targets (NES = −1.80, padj = 0.022) and Myc Targets V1 (NES = −1.70, padj = 0.022) alongside activation of Heme Metabolism (NES = +1.85, padj = 0.022) (**Fig. 4A**). Transcription factor activity analysis further highlighted sex-specific regulatory responses: PPARα was significantly activated in the colon of females (score = +3.40, padj = 0.042) but significantly suppressed in the liver of males (score = −3.63, padj = 0.022) (**Fig. 4B**), consistent with the opposing tissue directionality observed in the Hallmark analysis. Several biologically noteworthy examples of sex-specific regulation were identified at the protein level: Cyp2c11 was downregulated exclusively in the female liver (log₂FC = −1.84), Cyp2c6 was downregulated exclusively in the male liver (−1.46), and Fabp2 showed opposite-direction regulation in the colon (see above) (**Fig. S4B**). The antioxidant response was also sex-divergent, with females mounting a Gpx1/Gstp1-driven response in the liver while males showed reduced Nqo1 and Hmox1 (**Fig. S4C**). These findings emphasize the necessity of sex-stratified analysis in dietary intervention studies and suggest that the biological impact of DOxS is substantially modulated by hormonal or other sex-linked factors.

**Figure 4.**
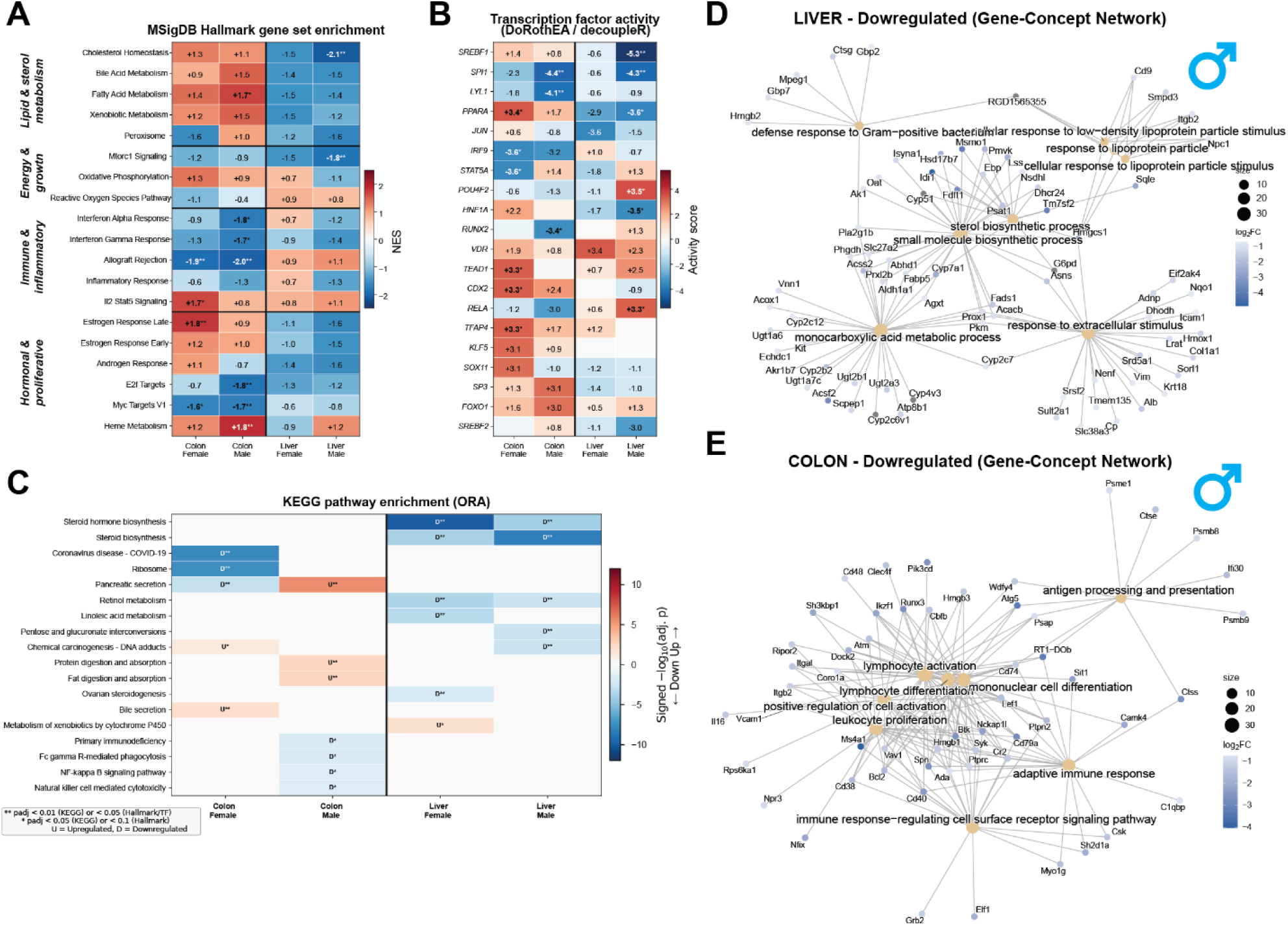
Pathway enrichment, transcription factor activity, and gene-concept network analyses reveal coordinated suppression of sterol metabolism in liver and immune signaling in colon. **(A)** MSigDB Hallmark gene set enrichment analysis (GSEA) across all four tissue × sex comparisons. Heatmap displays normalized enrichment scores (NES); asterisks indicate statistical significance (*padj < 0.1 and **padj < 0.05). Gene sets are grouped by functional theme. **(B)** Transcription factor activity inferred by decoupleR using the univariate linear model method with DoRothEA regulons (confidence levels A–C). Heatmap shows activity scores for the top transcription factors across comparisons. **(C)** KEGG pathway over-representation analysis (ORA) of upregulated (U) and downregulated (D) protein sets. Heatmap shows signed −log10(adjusted p-value), with red indicating enrichment among upregulated proteins and blue among downregulated. Significance thresholds: **p_adj < 0.01 or < 0.05; *p_adj < 0.05 or < 0.1. **(D)** Gene-concept network (cnetplot) of downregulated GO Biological Process terms in liver males, showing individual proteins mapped to enriched terms. Node color represents log2FC; node size reflects the number of associated terms. **(E)** Gene-concept network of downregulated GO Biological Process terms in colon males. Enrichment analyses used Benjamini-Hochberg FDR correction; DEPs were defined as |log2FC| > 0.585 and p < 0.05.

### Tissue Specificity and Shared Responses

The predominant finding across the dataset was the tissue specificity of DOxS-induced proteomic changes. Only 290 of 1,375 female colon DEPs were also differentially expressed in the female liver (21%), and only 224 of 1,149 male colon DEPs overlapped with the male liver (19%). This tissue divergence was strikingly reflected in the Hallmark analysis, where Cholesterol Homeostasis, Bile Acid Metabolism, Fatty Acid Metabolism, and Xenobiotic Metabolism showed opposing enrichment directions between tissues — positive NES in the colon and negative in the liver — revealing a mirror-image metabolic response to DOxS (**Fig. 4A**). KEGG pathway enrichment confirmed this dichotomy: the liver was dominated by suppression of steroid biosynthesis (padj = 6.2 × 10⁻L in males) and steroid hormone biosynthesis, while the colon showed suppression of immune pathways including NF-kappa B signaling, Fc gamma R-mediated phagocytosis, and natural killer cell mediated cytotoxicity (**Fig. 4C**). Gene-concept network analysis visualized the fundamentally distinct pathway architectures of each tissue: the hepatic network (**Fig. 4D**) was organized around sterol biosynthetic process and small molecule biosynthetic process, with hub genes Idi1, Fdft1, Cyp51, Msmo1, and Hmgcs1 bridging multiple metabolic processes, whereas the colonic network (**Fig. 4E**) was dominated by a dense immune cluster centered on lymphocyte activation, adaptive immune response, and leukocyte proliferation, with Btk, Syk, Ptprc, Cd74, and Ms4a1 as key hub genes. Upstream regulator analysis reinforced this tissue dichotomy: SREBF1 was the top suppressed transcription factor in the male liver (score = −5.32, padj = 2.5 × 10⁻L), whereas SPI1 (PU.1), a master regulator of myeloid/lymphoid differentiation, was the top suppressed TF in the male colon (score = −4.38, padj = 2.9 × 10⁻³) (**Fig. 4B**). One protein, alpha-1-antiproteinase (Serpina1), was the only named gene significantly downregulated across all four comparisons (colon female: log₂FC = −2.68; colon male: −2.43; liver female: −0.82; liver male: −0.72; all p < 0.05). Collectively, these results demonstrate that DOxS exert organ-specific effects – metabolic reprogramming in the liver and immune suppression in the colon – governed by distinct upstream regulators that cannot be extrapolated from one tissue to another.

### Metabolomics Reveals Tissue-Divergent Metabolic Responses to Dietary DOxS

To characterize DOxS-induced metabolic changes at the small-molecule level, we performed untargeted polar metabolomics on liver and colon tissues using timsTOF. Metabolite annotation employed a multi-criteria confidence framework (MS/MS matching, mSigma, Δm/z < 5 ppm, ΔCCS < 5%), yielding 176 high-confidence metabolites in the liver and 104 in the colon (**Fig. 5A-B**). Differential abundance was assessed using Mann-Whitney U tests with Benjamini-Hochberg correction (significance: padj < 0.05, |FC| > 1.5). Ten metabolites reached significance in the liver and 21 in the colon, with only one shared between tissues. The dominant hepatic signal was broad phospholipid accumulation under DOxS (**Fig. 5C**). Seven of 10 significant metabolites were lipids elevated in the DOxS group, including LPA(18:1) (log2FC = +2.97), LPC(14:0) (+1.69), PG(18:2/18:2) (+10.68), and PE(16:0/20:4) (+2.30), all padj < 0.03.

**Figure 5.**
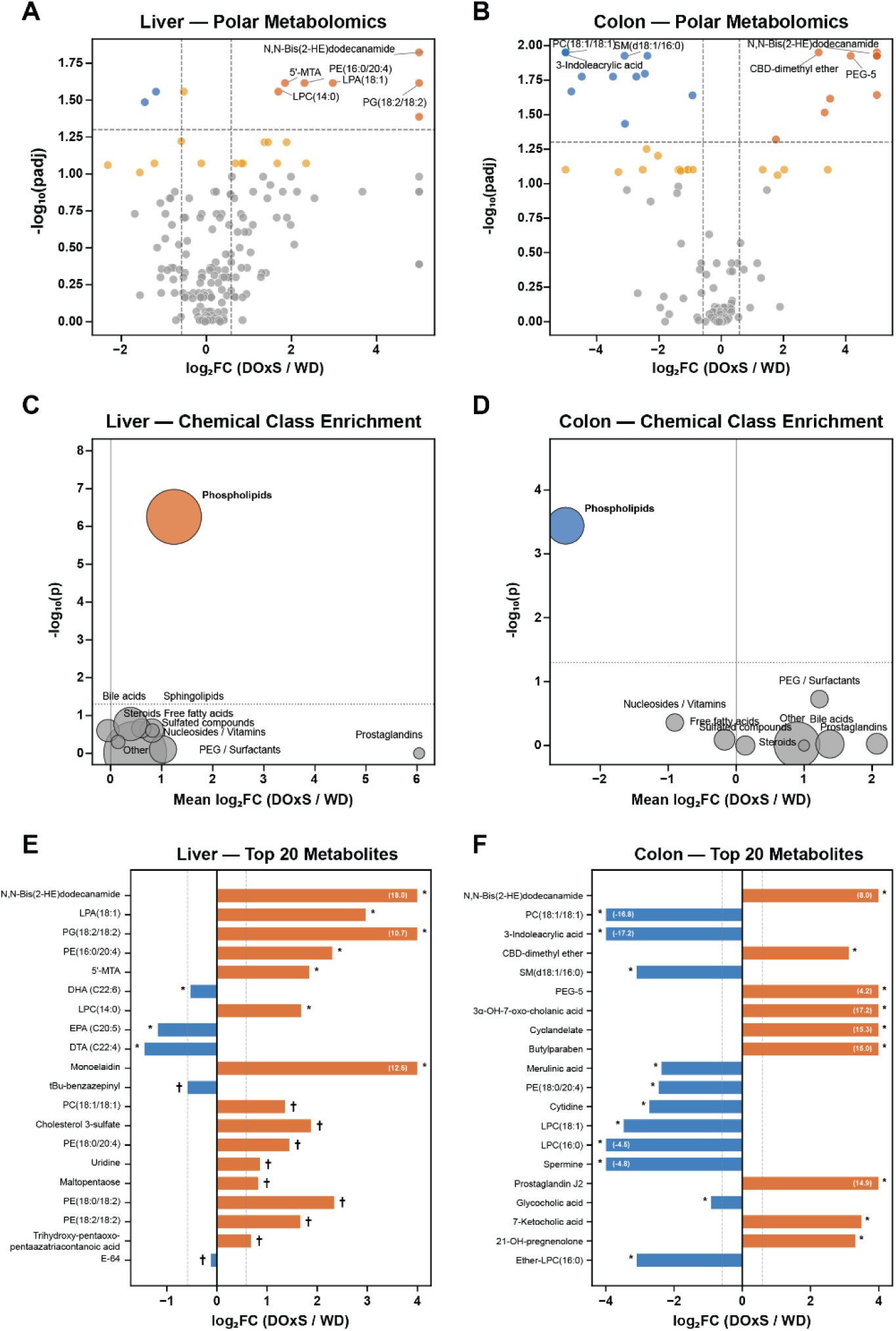
Polar metabolomics reveals tissue-divergent metabolic responses to dietary DOxS in liver and colon. **(A,B)** Volcano plots of liver and colon polar metabolites. Each point represents a single metabolite; red, significantly elevated in WD-DOxS (padj < 0.05, |log₂FC| > 0.585); blue, significantly elevated in WD; orange, trending (padj < 0.10); grey, not significant. Dashed lines indicate significance thresholds. **(C,D)** Chemical class enrichment analysis. Each bubble represents a metabolite class; bubble size is proportional to the number of annotated metabolites; color indicates directionality of the significant class (red, elevated in WD-DOxS; blue, elevated in WD; grey, not significant by Wilcoxon signed-rank test). Dotted line, p = 0.05. **(E,F)** Log₂ fold change for the 20 most differentially abundant metabolites in each tissue, ranked by raw p-value. **padj < 0.01; *padj < 0.05; †padj < 0.10. Differential abundance was assessed by two-sided Mann-Whitney U tests with Benjamini-Hochberg correction.

Across all 49 phospholipid annotations, 40 of 49 showed a positive direction (class mean log2FC = +1.24, Wilcoxon P < 0.001), providing metabolite-level confirmation of the Scd1/Fasn/Plin2 de novo lipogenesis program identified in the proteome. Conversely, the anti-inflammatory PUFAs EPA (C20:5; −1.18), DTA (C22:4; −1.45), and DHA (C22:6; −0.53) were all significantly depleted, consistent with Scd1-mediated competition for shared desaturase machinery and consumption of free PUFAs as acyl donors.^81^ 5′-Methylthioadenosine (MTA), a methionine salvage intermediate, was elevated 3.6-fold (padj = 0.024), converging with the metagenomic enrichment of methionine/SAM/THF biosynthesis in the DOxS gut microbiome. Notably, none of 18 bile acid annotations reached significance in the liver, consistent with Cyp7a1 suppression^82^ (4–5.5-fold in the proteome) shutting down hepatic bile acid synthesis.

The colon exhibited a phospholipid signature diametrically opposed to the liver. Six significant hits were phospholipids depleted under DOxS, including LPC(16:0) (−4.48), LPC(18:1) (−3.48), sphingomyelin (−3.10), and PC(18:1/18:1) (absent in all DOxS colons). The class mean log2FC was −2.51 across 22 annotations (Wilcoxon P < 0.001). This tissue-divergent phospholipid response—accumulation in liver, depletion in colon—is consistent with the liver upregulating de novo lipogenesis while the colon is deprived of biliary lipid delivery due to Cyp7a1 suppression.^82^ Unlike the liver, the colon showed significant bile acid remodeling. 3α-Hydroxy-7-oxo-5β-cholanic acid, an oxidized bile acid, was absent from all WD samples but present in 7 of 9 DOxS colons (padj = 0.012), and 7-ketocholic acid was elevated 11-fold (padj = 0.024), while glycocholic acid was reduced (padj = 0.023) (**Fig. 5E-F**). This shift from conjugated primary toward oxidized keto bile acids is consistent with the WD-DOxS-enriched bile acid-transforming taxa (*Alistipes*, *Parabacteroides*, *Ruminococcus gnavus*) identified in the metagenome.^83, 84^ Three colon-specific findings establish direct links to the other omics layers. First, prostaglandin J2 ^85^, an anti-inflammatory PPARγ ligand and JAK-STAT inhibitor,^86^ appeared exclusively in DOxS colons (6/9 DOxS vs. 0/10 WD; padj = 0.023), providing a candidate lipid mediator for the Irf9–ISG uncoupling paradox identified in the proteome. Second, 3-indoleacrylic acid, a microbial tryptophan metabolite and aryl hydrocarbon receptor (AhR) agonist critical for barrier integrity,^87^ was present in 9/10 WD colons but completely absent from all DOxS colons (padj = 0.011), directly traceable to the loss of its producer *Limosilactobacillus reuteri*^88^, the sole WD-enriched species in the metagenome. Third, spermine was depleted ∼28-fold (padj = 0.022), which, paired with elevated hepatic MTA, indicates a cross-tissue disruption of the SAM–polyamine axis spanning microbial production, hepatic metabolism, and colonic utilization.^89^

## DISCUSSION

Dietary oxysterols^13^ are among the most abundant and biologically active oxidized lipids in Western diets, yet their systemic effects beyond acute cytotoxicity remain poorly understood.^13, 15, 18^ By integrating shotgun metagenomics, transcriptomics, quantitative proteomics, and untargeted metabolomics across two target tissues and the gut microbiome, the present study provides the first multi-omics portrait of how DOxS exposure reshapes host metabolism when superimposed on a Western diet. Three convergent themes emerge from this integration: (i) a near-complete shutdown of hepatic cholesterol biosynthesis coupled with a paradoxical activation of de novo lipogenesis, (ii) a profound remodeling of the gut microbiota with functionally consequential loss of key commensals, and (iii) a striking tissue specificity in which liver and colon frequently mount divergent or even opposing metabolic and immune responses to the same dietary challenge.

The most prominent hepatic response to DOxS exposure was the near-complete suppression of the mevalonate pathway, evident from the coordinated downregulation of virtually every enzyme from HMGCS1 through FDFT1 (**Fig. 3**). This finding aligns with the canonical mechanism by which oxysterols regulate cholesterol homeostasis: binding to SCAP–Insig complexes in the endoplasmic reticulum retains SREBP-2 in an inactive state, thereby silencing its transcriptional targets throughout the cholesterol biosynthetic cascade.^21, 23, 75^ Importantly, the effect in our study was sex-dependent, with male livers exhibiting a more profound suppression than females (**Fig. 3D**). This observation is consistent with the known sexual dimorphism in hepatic sterol metabolism^90^ and suggests that DOxS may interact with sex-hormone-regulated pathways governing SREBP-2 activity. Transcription-factor activity inference confirmed SREBF1 as the most suppressed regulator in male livers, corroborating the proteomic data at the regulatory level.

Paradoxically, despite SREBF1 suppression, DOxS-treated livers showed marked upregulation of the de novo lipogenesis (DNL) enzymes Scd1, Fasn, and Plin2 (**Fig. 3E**). This apparent contradiction can be reconciled through the dual signaling nature of oxysterols: while they suppress SREBP processing via Insig,^75^ they simultaneously serve as potent ligands for liver X receptors (LXRs), which directly transactivate *Srebf1c*, *Fasn*, and *Scd1* independently of SREBP-2.^22, 91^ Indeed, LXR activation by oxysterols has been shown to drive hepatic steatosis in vivo through SREBP-1c-mediated lipogenesis even when cholesterol synthesis is suppressed.^92^ Our metabolomics data reinforce this model: phospholipid species accumulated in DOxS livers while polyunsaturated fatty acids (PUFAs) were depleted, a pattern consistent with increased Scd1 activity converting saturated substrates to monounsaturated products for phospholipid remodeling (**Fig. 5**). These findings extend prior work by our group documenting the widespread occurrence and pro-inflammatory potential of DOxS in the food supply,^13, 15, 28, 93^ and demonstrate that chronic dietary exposure at physiologically relevant levels is sufficient to reprogram hepatic lipid metabolism in vivo.

A striking observation is the near-complete disconnect between the transcriptomic and proteomic layers. RNA-seq across liver, heart, and brain yielded only a handful of differentially expressed genes in response to DOxS treatment – Rfx6 and Sephs2 in liver (both restricted to females), Mug1 and Apoa4 in brain, and Moxd1, Cr2, and Cd19 in heart (**Fig. S2**) – while the liver proteome revealed thousands of quantitative changes converging on clearly defined metabolic pathways (**Fig. 3**). This discordance may reflect the fundamental biology of oxysterol signaling. The SREBP pathway operates primarily through regulated intramembrane proteolysis: oxysterols prevent the SCAP–SREBP complex from transiting to the Golgi, blocking the proteolytic release of mature SREBPs without necessarily altering SREBP mRNA levels.^21, 94^ Similarly, oxysterol-mediated degradation of HMGCR proceeds through Insig-dependent ubiquitination and proteasomal turnover, a post-translational mechanism invisible to RNA-seq.^95^ LXR target gene induction (e.g., Fasn, Scd1) might be expected at the transcript level, but could be masked in bulk tissue by cellular heterogeneity or by compensatory transcript degradation. The transcriptome–proteome disconnect thus underscores that dietary oxysterols act predominantly through post-transcriptional and post-translational mechanisms, making proteomics and metabolomics the appropriate discovery platforms for this class of dietary bioactive compounds.

Among the sparse transcriptomic hits, the downregulation of Apoa4 in DOxS-treated brains is noteworthy. ApoA-IV is a lipid transport apolipoprotein with documented roles in central lipid sensing and neuroprotection,^96^ and its suppression may reflect altered cholesterol trafficking in the CNS under oxysterol pressure. In the liver, the sex-specific induction of Rfx6—a transcription factor involved in endocrine cell differentiation—hints at potential metabolic-endocrine crosstalk in females, though the biological significance of this isolated finding remains uncertain. The cardiac hits (Cr2, Cd19) suggest possible B-cell or complement-related immune activation but were driven by only a few samples and should be interpreted cautiously. Collectively, the transcriptomic layer serves as a valuable negative control, reinforcing the conclusion that oxysterol-driven metabolic reprogramming operates below the transcriptional radar.

Convergent evidence from proteomics and metabolomics demonstrated suppression of the classical bile acid synthesis pathway. Cyp7a1, the rate-limiting enzyme, was markedly reduced at the protein level in DOxS livers, and this was mirrored by decreased abundance of primary bile acid intermediates in liver metabolomics. In the colon, however, bile acid composition shifted rather than uniformly declined, with certain secondary bile acids appearing preferentially in DOxS-treated animals. This tissue-level divergence suggests that while hepatic bile acid production is curtailed—likely via FXR-mediated feedback amplified by the oxysterol burden—the colonic microbiota continues to actively transform the reduced bile acid pool, potentially generating species with distinct signaling properties.^64, 83^ The concurrent suppression of Cyp7a1 and the mevalonate pathway reinforces the interpretation that DOxS act as potent endocrine disruptors of cholesterol catabolism, blocking both its synthesis and its major disposal route.

Dietary DOxS profoundly remodeled the gut microbiome, increasing α-diversity and driving compositional separation detectable by PERMANOVA (**Fig. 2**). Of the 20 species identified as differentially abundant by MaAsLin2, 13 were enriched in the WD-DOxS group, whereas the control Western diet was characterized by the dominance of a small number of taxa, notably *Ligilactobacillus murinus* and *Limosilactobacillus reuteri*. The depletion of *L. reuteri* is of particular functional significance because this species is a primary producer of tryptophan-derived aryl hydrocarbon receptor (AhR) agonists, including indole-3-lactic acid and indoleacrylic acid, through the aromatic amino acid aminotransferase (ArAT) pathway.^66, 88, 97^ Strikingly, 3-indoleacrylic acid (IA) was completely absent in the colonic metabolome of DOxS-treated animals. IA has been shown to promote intestinal epithelial barrier integrity and suppress inflammatory responses via AhR activation in both epithelial cells and macrophages.^87,98^ Its complete loss in DOxS-treated colons, coupled with the depletion of its known bacterial producers, establishes a microbiome–metabolite axis linking dietary oxysterols to impaired mucosal defense.

A defining feature of this study is the tissue specificity of metabolic responses. Phospholipid species that accumulated in DOxS-treated livers were depleted in DOxS-treated colons, suggesting an organ-level redistribution rather than a systemic increase (**Fig. 5**). This “phospholipid flip” may reflect the hepatic diversion of lipid substrates toward storage and VLDL assembly under LXR activation, while the colon, deprived of normal bile acid-mediated lipid delivery and simultaneously losing barrier-protective microbial metabolites, undergoes phospholipid depletion. Tissue-resolved analyses are therefore essential for understanding DOxS toxicology: pooling liver and colon data would mask these opposing signals and lead to erroneous conclusions about the net metabolic impact of dietary oxysterols.

The colonic proteome revealed a distinctive immune-signaling pattern characterized by IRF9 upregulation without proportional induction of canonical interferon-stimulated genes ^77^. This IRF9–ISG uncoupling is unusual, as IRF9 typically forms the ISGF3 complex with STAT1/STAT2 to drive ISG transcription (**Fig. 4**).^78^ One interpretation is that DOxS-derived signals activate IRF9 via non-canonical pathways—possibly through unfolded protein response cross-talk or lipid-mediated NF-κB modulation—without fully engaging the type I interferon cascade. Supporting this, prostaglandin J2^85^ appeared exclusively in DOxS-treated colons. PGJ2 and its derivative 15-deoxy-Δ12,14-PGJ2 are potent anti-inflammatory lipid mediators that inhibit NF-κB signaling through both PPARγ-dependent and -independent mechanisms, including covalent modification of IκB kinase.^85, 86^ The exclusive presence of PGJ2 in DOxS colons may represent a compensatory anti-inflammatory response to the pro-oxidant burden of dietary oxysterols, operating in parallel with the IRF9 pathway to modulate colonic immune tone. Notably, the pro-inflammatory and pro-apoptotic activities of DOxS have been extensively documented in cellular models,^15^ and the colonic responses observed here likely reflect the tissue’s attempt to counterbalance these effects in vivo.^19^

Cross-omics integration also revealed a methionine–polyamine axis connecting the gut microbiome to hepatic one-carbon metabolism. 5′-Methylthioadenosine (MTA), a byproduct of polyamine biosynthesis, was elevated in DOxS livers, while spermine was depleted in colonic tissue (**Fig. 5**). This pattern is consistent with accelerated polyamine flux in the liver, possibly driven by ornithine decarboxylase activation under cellular stress, and impaired polyamine recycling or uptake in the colon. Given that several of the DOxS-enriched microbial taxa are known participants in methionine salvage pathways, the microbiome may directly contribute to the altered MTA–spermine balance. Polyamine depletion in the colon has been linked to impaired epithelial proliferation and barrier dysfunction,^99, 100^ providing yet another route through which dietary oxysterols may compromise intestinal homeostasis.

In summary, this integrated multi-omics investigation reveals that dietary DOxS, at levels consistent with Western diet exposure, orchestrate a coordinated reprogramming of hepatic lipid metabolism—simultaneously shutting down cholesterol biosynthesis while paradoxically activating de novo lipogenesis—coupled with a profound restructuring of the gut microbiome that eliminates key barrier-protective commensals and their tryptophan-derived metabolites. The near-silence of the hepatic transcriptome in the face of sweeping proteomic changes establishes that oxysterols operate predominantly through post-transcriptional mechanisms, a finding with important implications for study design in nutritional toxicology. The tissue-specific and often opposing metabolic responses in liver and colon underscore the importance of organ-resolved analyses. These findings position dietary oxysterols not merely as markers of food processing quality, but as biologically active modulators of the gut–liver axis with implications for metabolic disease, intestinal barrier integrity, and immune regulation. Future studies incorporating dose–response designs, longitudinal sampling, and integration of the lipidomics layer will be essential to establish the translational relevance of these findings to human dietary exposures.

### Limitations of the study

First, the study employed a single DOxS concentration superimposed on one dietary background (Total Western Diet) in Wistar rats, precluding dose–response characterization and limiting direct extrapolation to human exposures, which span a wide range across food categories.^13^ Future studies should incorporate graded DOxS doses informed by human exposure estimates from the FooDOxS database, ideally in multiple rodent strains or humanized microbiota models to improve translational relevance. Second, the current analysis lacks a dedicated lipidomics layer and did not include colonic tissue in the RNA-seq design due to RNA degradation.^73^ Integrating targeted lipidomics — particularly oxysterol, bile acid, and phospholipid panels — across both tissues would strengthen the mechanistic links inferred here between Scd1 upregulation, phospholipid redistribution, and bile acid suppression, and would enable direct quantification of the DOxS species reaching each tissue compartment.

## Acknowledgments

We thank CosmosID for support in analyzing and interpreting the shotgun metagenomic data, and Jennifer Hackett at the University of Kansas Genome Sequencing Core for sequencing services. This study was partially funded by USDA grant 2021-67012-35038 to L.M.P. and I.G.M.M., and by start-up funding from the University of Kansas and the University of Kansas Cancer Center (KUCC) to C.B. The KU Genome Sequencing Core is supported by the National Institute of General Medical Sciences of the National Institutes of Health under Award Number P30 GM145499.

## Author contributions (CRediT)

Lisaura Maldonado-Pereira: *Investigation (animal intervention, pharmacokinetics, DOxS analysis), Methodology, Data curation, Writing – original draft*. Thuraya M. Mutawi: *Investigation (RNA extraction, RNA-seq library preparation), Formal analysis.* Arunima Singh: *Investigation (RNA extraction)*. Brian J. Sanderson: *Formal analysis (RNA-seq computational analysis), Software, Visualization (RNA-seq figures), Writing – review & editing.* Michaella J. Rekowski: *Resources (mass spectrometry core facility), Supervision (mass spectrometry)*. Carlo Barnaba: *Conceptualization, Methodology, Formal analysis (proteomics, metagenomics), Funding, Software, Visualization, Writing – original draft, Writing – review & editing, Supervision, Funding acquisition, Project administration*. Ilce G. Medina-Meza: *Conceptualization, Funding, Methodology, Formal analysis (metabolomics), Investigation, Writing – original draft, Writing – review & editing, Supervision, Funding acquisition, Project administration*.

---

**Figure S1.**
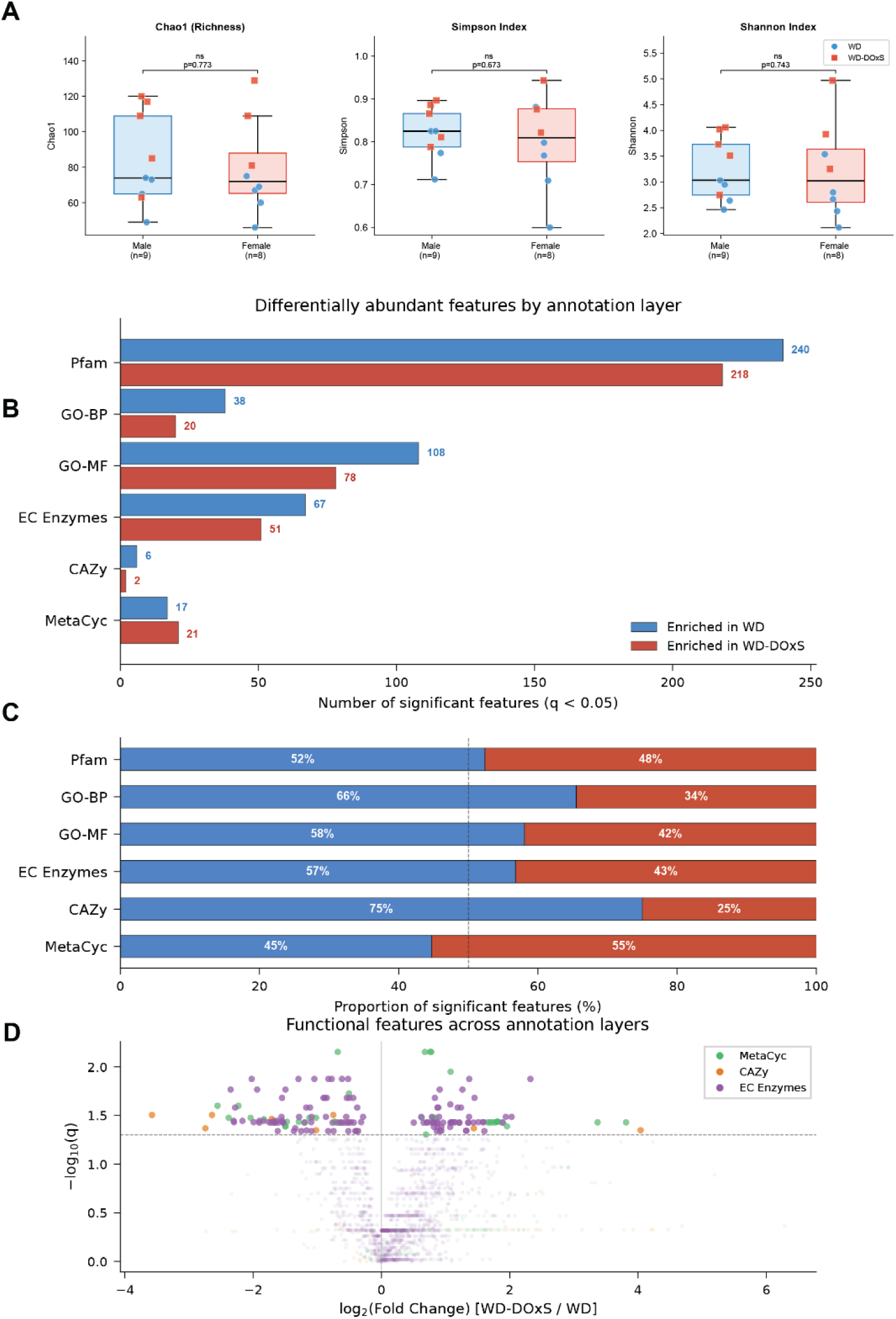
Sex does not confound diet-driven microbiome changes and functional metagenomics reveal asymmetric enrichment patterns across annotation layers. **(A)** Alpha diversity (Chao1 richness, Simpson index, Shannon index) compared between males and females, pooled across diets. Boxes represent the interquartile range with the median indicated by a horizontal line; whiskers extend to 1.5 × IQR. Individual samples are overlaid as points colored by diet (blue circles, WD; red squares, WD-DOxS). No significant sex effect was detected for any metric (Mann-Whitney U test, all *P* > 0.67). **(B)** Number of differentially abundant features (q < 0.05, Mann–Whitney U with Benjamini–Hochberg correction) enriched in WD (blue) versus WD-DOxS (red) across six functional annotation layers from CosmosID: Pfam, GO-BP (biological process), GO-MF (molecular function), EC Enzymes, CAZy (carbohydrate-active enzymes), and MetaCyc metabolic pathways. Feature counts are displayed at bar termini. Pfam yielded the largest number of significant features in both directions (240 WD, 218 WD-DOxS), while CAZy was the sparsest (6 WD, 2 WD-DOxS). **(C)** Proportional representation of significant features enriched in each diet group. WD dominated in most layers (52–75%), with the notable exception of MetaCyc, where WD-DOxS accounted for 55% of significant pathways. Dashed line indicates the 50% midpoint. **(D)** Combined volcano plot of functional features across three enzyme/pathway annotation layers (MetaCyc, green; CAZy, orange; EC Enzymes, purple). The x-axis shows log₂ fold change (WD-DOxS/WD; positive values indicate WD-DOxS enrichment), and the y-axis shows –log₁₀(q). Dashed horizontal line indicates q = 0.05 significance threshold.

**Figure S2.**
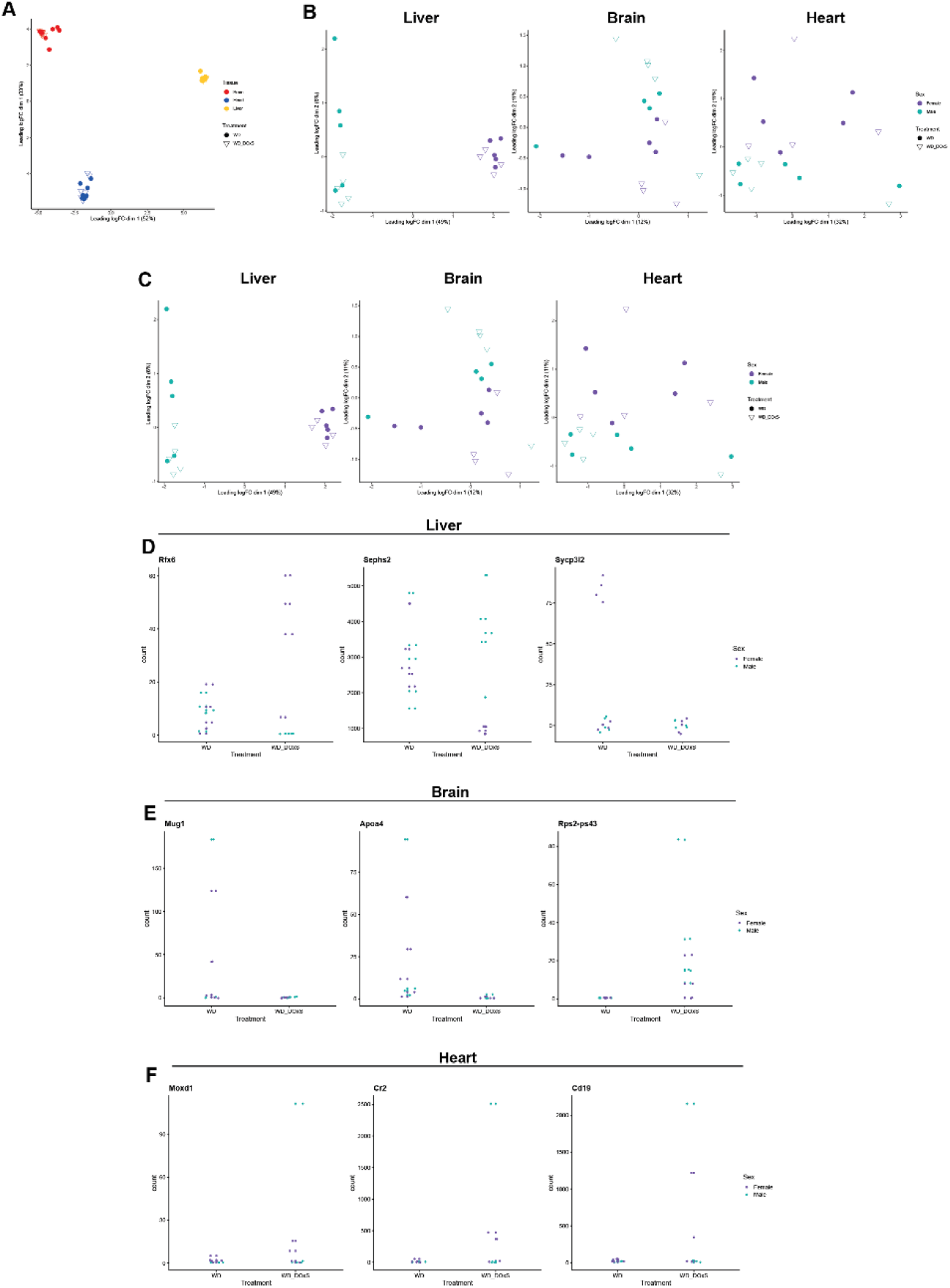
Transcriptomic profiling of liver, heart, and brain reveals minimal differential gene expression in response to dietary DOxS. **(A)** Multidimensional scaling (MDS) plot of raw gene expression counts across all samples, showing clear clustering by tissue. Tissues are color-coded (brain, red; heart, teal; liver, blue); treatment is indicated by shape (filled circles, WD; open triangles, WD-DOxS). **(B)** Within-tissue MDS plots (liver, brain, heart) prior to outlier removal, with samples colored by sex (female, purple; male, green) and shaped by treatment (filled circles, WD; open triangles, WD-DOxS). **(C)** Within-tissue MDS plots after exclusion of the brain outlier, showing modest separation by sex and treatment across all three tissues. **(D)** Liver: strip plots of raw expression counts for genes with significant sex-by-treatment interaction (*Rfx6*, *Sephs2*) or main effect of DOxS (*Sycp3l2*). Sex indicated by color (purple, female; green, male). **(E)** Brain: strip plots for *Mug1* and *Apoa4* (both decreased by DOxS) and *Rps2-ps43* (increased). (F) Heart: strip plots for *Moxd1*, *Cr2*, and *Cd19*. For panels D–F, individual data points are colored by sex (purple, female; green, male). Statistical models and full results are reported in Tables S3–S4.

**Fig. S3.**
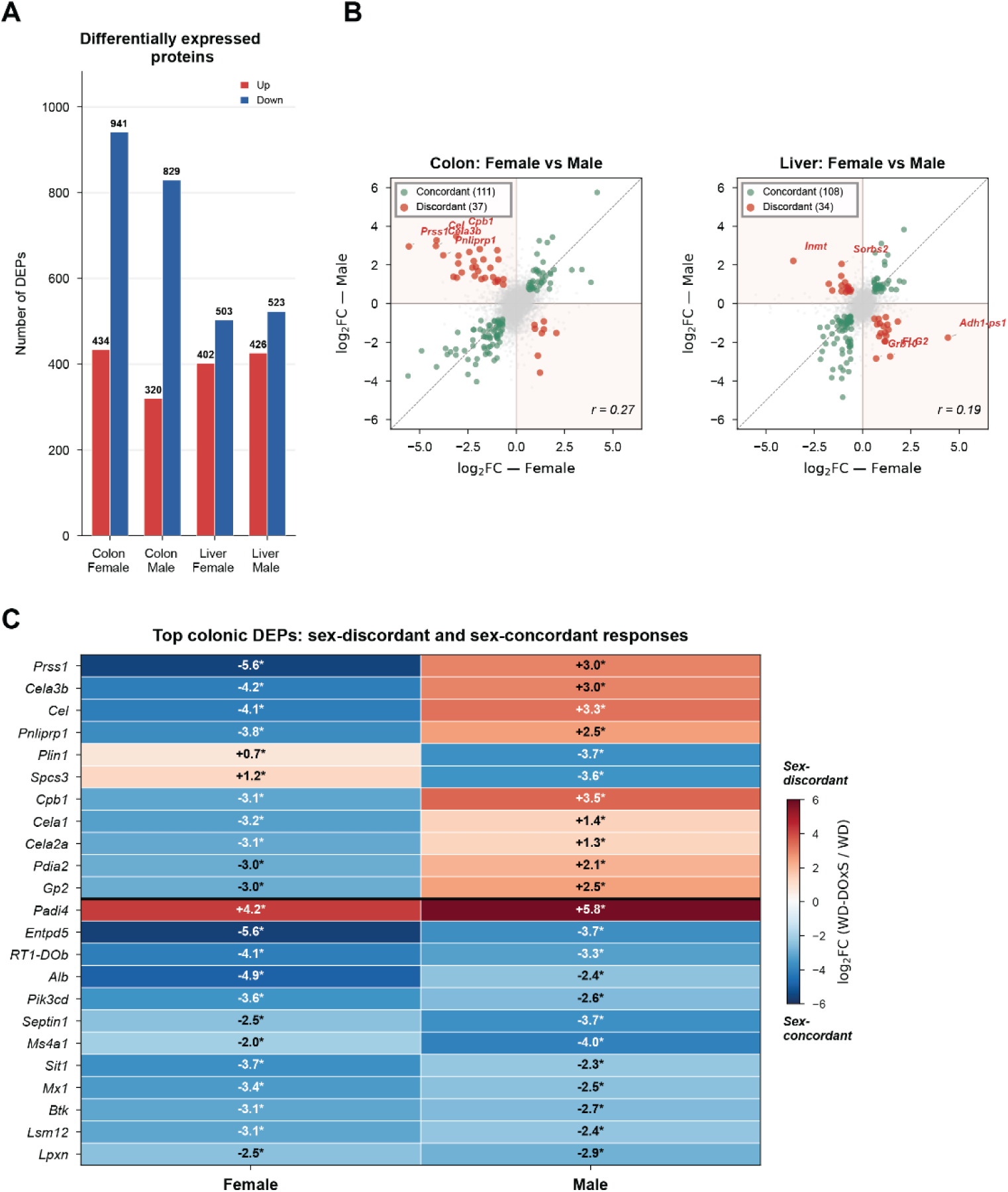
Global proteomics landscape and sex dimorphism in the proteomic response to dietary DOxS. **(A)** Number of differentially expressed proteins (DEPs; |log₂FC| > 0.585, p < 0.05) identified in each tissue × sex comparison (WD-DOxS vs WD). Red, upregulated; blue, downregulated. **(B)** Scatter plots of log₂FC values in females vs males for all shared quantified proteins in the colon (left) and liver (right). Each dot represents one protein. Green dots indicate DEPs significant in both sexes with concordant direction; red dots indicate significant DEPs with discordant direction (opposite fold-change between sexes). Shaded quadrants highlight discordant zones. Pearson correlation coefficients are shown. Key discordant genes are labeled. **(C)** Heatmap of top colonic DEPs exhibiting sex-discordant (above black line) or sex-concordant (below) responses. Values indicate log₂FC; asterisks denote significance (p < 0.05, |log₂FC| > 0.585). Sex-discordant proteins were predominantly digestive enzymes (Prss1, Cela3b, Cel, Cpb1), while sex-concordant proteins included immune markers (RT1-DOb, Ms4a1, Mx1, Btk) suppressed in both sexes.

**Fig. S4.**
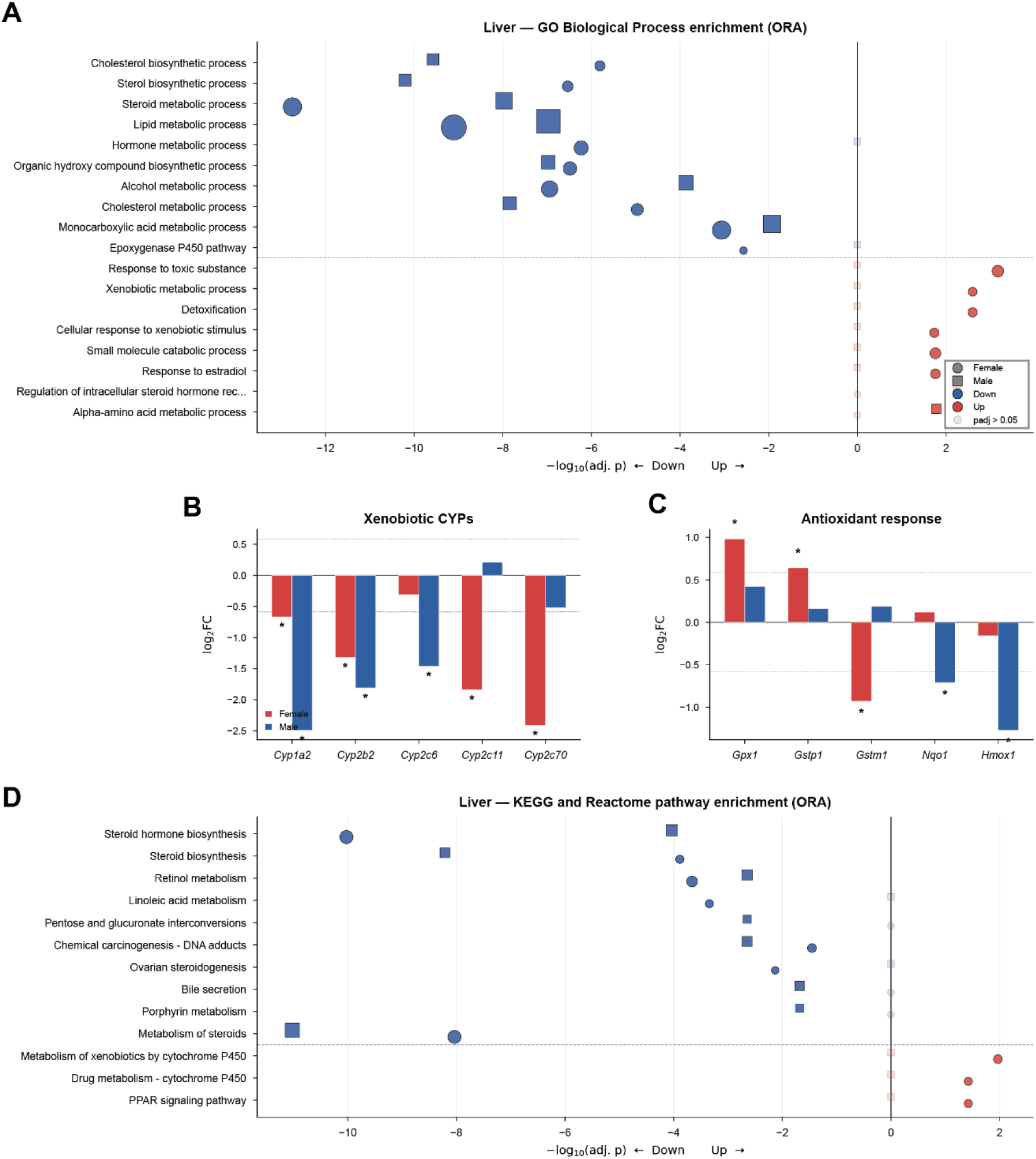
Extended hepatic pathway and protein-level analysis. **(A)** Bidirectional dot plot of Gene Ontology Biological Process (GO-BP) enrichment in the liver. Circles, females; squares, males. Dot size is proportional to gene count; opacity indicates significance (filled, padj < 0.05; faded, padj > 0.05). Left of the vertical line represents downregulated pathways; right represents upregulated. **(B)** Log₂FC of hepatic xenobiotic-metabolizing cytochrome P450 enzymes. **(C)** Log₂FC of hepatic antioxidant response proteins. **(D)** Bidirectional dot plot of KEGG and Reactome pathway enrichment in the liver. Asterisks in panels **B** and **C** denote *p* < 0.05 and |log₂FC| > 0.585. Dashed lines in **B** and **C** indicate the ±0.585 fold-change threshold

**Fig. S5.**
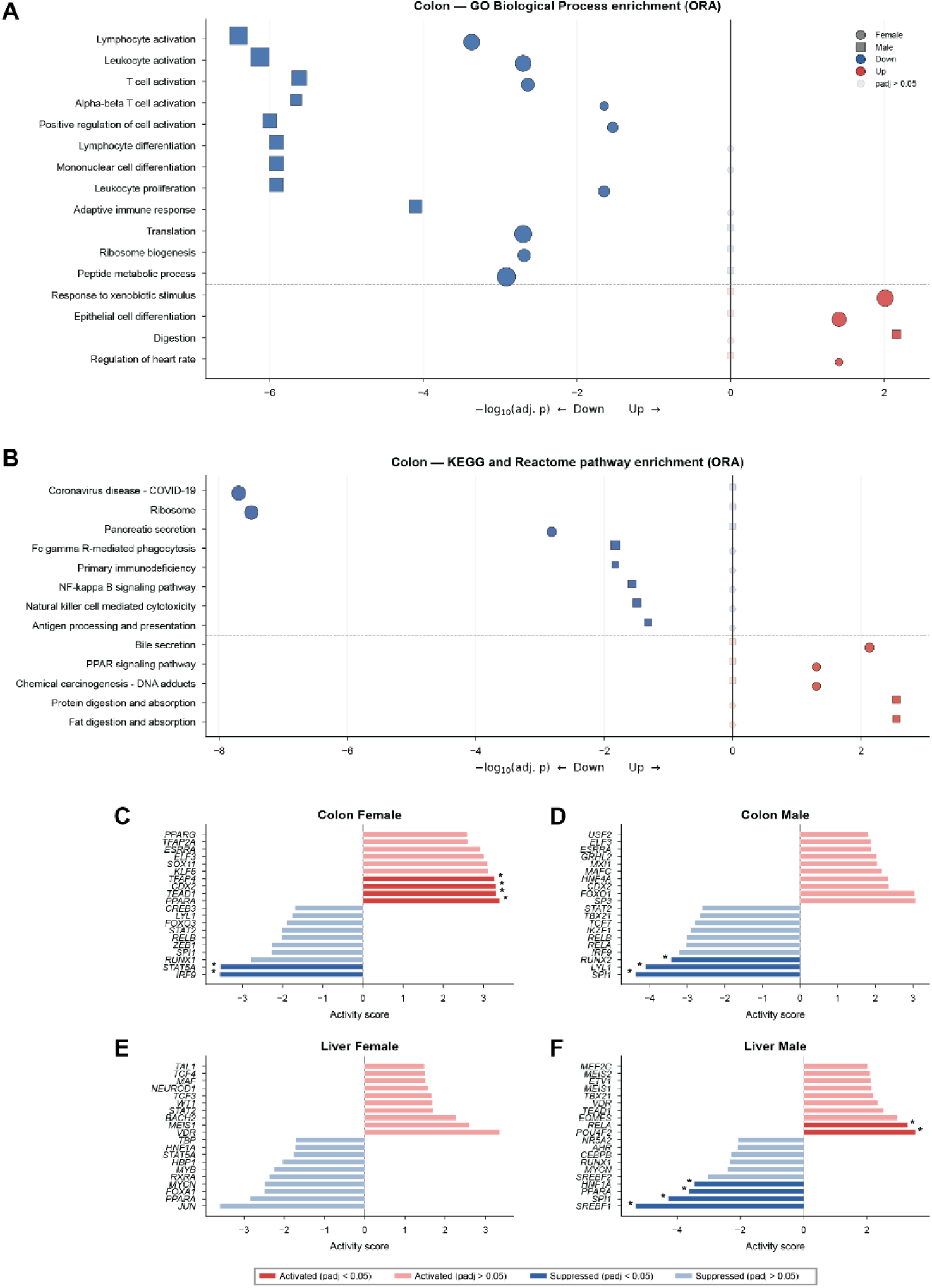
Extended colonic pathway analysis and transcription factor activity across all comparisons. **(A)** Bidirectional dot plot of GO-BP enrichment in the colon. **(B)** KEGG and Reactome pathway enrichment in the colon. **(C-F)** Transcription factor (TF) activity scores inferred by decoupleR using the univariate linear model (ULM) method with DoRothEA regulons (confidence A–C; 271 TFs, 13,223 interactions). Bars show the top 10 activated and top 10 suppressed TFs for each comparison ranked by absolute activity score. Dark bars indicate padj < 0.05; pale bars indicate padj > 0.05.

## REFERENCES

1. Monteiro CA, Cannon G, Levy RB, Moubarac JC, Louzada ML, Rauber F, Khandpur N, Cediel G, Neri D, Martinez-Steele E, Baraldi LG, Jaime PC. Ultra-processed foods: what they are and how to identify them. Public Health Nutr. 2019;22(5):936–41. Epub 20190212. doi: 10.1017/S1368980018003762. PubMed PMID: 30744710; PMCID: PMC10260459.

2. Menichetti G, Ravandi B, Mozaffarian D, Barabasi AL. Machine learning prediction of the degree of food processing. Nat Commun. 2023;14(1):2312. Epub 20230421. doi: 10.1038/s41467-023-37457-1. PubMed PMID: 37085506; PMCID: PMC10121643.

3. van Boekel M, Fogliano V, Pellegrini N, Stanton C, Scholz G, Lalljie S, Somoza V, Knorr D, Jasti PR, Eisenbrand G. A review on the beneficial aspects of food processing. Mol Nutr Food Res. 2010;54(9):1215–47. doi: 10.1002/mnfr.200900608. PubMed PMID: 20725924.

4. Wahlqvist ML. Food structure is critical for optimal health. Food Funct. 2016;7(3):1245–50. doi: 10.1039/c5fo01285f. PubMed PMID: 26667120.

5. Zobel EH, Hansen TW, Rossing P, von Scholten BJ. Global Changes in Food Supply and the Obesity Epidemic. Curr Obes Rep. 2016;5(4):449–55. doi: 10.1007/s13679-016-0233-8. PubMed PMID: 27696237.

6. Adolph TE, Tilg H. Western diets and chronic diseases. Nature Medicine. 2024;30(8):2133–47.

7. Fitzpatrick JA, Halmos EP, Gibson PR, Machado PP. Ultra-processed Foods and Risk of Crohn’s Disease: How Much is Too Much? Clin Gastroenterol Hepatol. 2023;21(10):2478–80. Epub 20230324. doi: 10.1016/j.cgh.2023.03.009. PubMed PMID: 36967099.

8. Spreadbury I. Comparison with ancestral diets suggests dense acellular carbohydrates promote an inflammatory microbiota, and may be the primary dietary cause of leptin resistance and obesity. Diabetes Metab Syndr Obes. 2012;5:175–89. Epub 20120706. doi: 10.2147/DMSO.S33473. PubMed PMID: 22826636; PMCID: PMC3402009.

9. Grundy MM, Lapsley K, Ellis PR. A review of the impact of processing on nutrient bioaccessibility and digestion of almonds. Int J Food Sci Technol. 2016;51(9):1937–46. Epub 20160731. doi: 10.1111/ijfs.13192. PubMed PMID: 27642234; PMCID: PMC5003169.

10. Arnone D, Vallier M, Hergalant S, Chabot C, Ndiaye NC, Moulin D, Aignatoaei AM, Alberto JM, Louis H, Boulard O, Mayeur C, Dreumont N, Peuker K, Strigli A, Zeissig S, Hansmannel F, Chamaillard M, Kokten T, Peyrin-Biroulet L. Long-Term Overconsumption of Fat and Sugar Causes a Partially Reversible Pre-inflammatory Bowel Disease State. Front Nutr. 2021;8:758518. Epub 20211118. doi: 10.3389/fnut.2021.758518. PubMed PMID: 34869528; PMCID: PMC8637418.

11. Aguayo-Patron SV, Calderon de la Barca AM. Old Fashioned vs. Ultra-Processed-Based Current Diets: Possible Implication in the Increased Susceptibility to Type 1 Diabetes and Celiac Disease in Childhood. Foods. 2017;6(11). Epub 20171115. doi: 10.3390/foods6110100. PubMed PMID: 29140275; PMCID: PMC5704144.

12. Dahl WJ, Rivero Mendoza D, Lambert JM. Diet, nutrients and the microbiome. Prog Mol Biol Transl Sci. 2020;171:237–63. Epub 20200425. doi: 10.1016/bs.pmbts.2020.04.006. PubMed PMID: 32475524.

13. Medina-Meza IG, Vaidya Y, Barnaba C. FooDOxS: a database of oxidized sterols content in foods. Food Funct. 2024;15(12):6324–34. Epub 20240617. doi: 10.1039/d4fo00678j. PubMed PMID: 38726678.

14. Maldonado-Pereira L, Barnaba C, Medina-Meza IG. Oxidative Status of Ultra-Processed Foods in the Western Diet. Nutrients. 2023;15(23). Epub 20231122. doi: 10.3390/nu15234873. PubMed PMID: 38068731; PMCID: PMC10708126.

15. Maldonado-Pereira L, Schweiss M, Barnaba C, Medina-Meza IG. The role of cholesterol oxidation products in food toxicity. Food Chem Toxicol. 2018;118:908–39. Epub 20180627. doi: 10.1016/j.fct.2018.05.059. PubMed PMID: 29940280.

16. Savage GP, Dutta PC, Rodriguez-Estrada MT. Cholesterol oxides: their occurrence and methods to prevent their generation in foods. Asia Pac J Clin Nutr. 2002;11(1):72–8. doi: 10.1046/j.1440-6047.2002.00270.x. PubMed PMID: 11890642.

17. Lemaire S, Lizard G, Monier S, Miguet C, Gueldry S, Volot F, Gambert P, Néel D. Different patterns of IL-1β secretion, adhesion molecule expression and apoptosis induction in human endothelial cells treated with 7α-, 7β-hydroxycholesterol, or 7-ketocholesterol. FEBS letters. 1998;440(3):434–9.

18. Testa G, Rossin D, Poli G, Biasi F, Leonarduzzi G. Implication of oxysterols in chronic inflammatory human diseases. Biochimie. 2018;153:220–31. Epub 20180609. doi: 10.1016/j.biochi.2018.06.006. PubMed PMID: 29894701.

19. Willinger T. Oxysterols in intestinal immunity and inflammation. J Intern Med. 2019;285(4):367–80. Epub 20181126. doi: 10.1111/joim.12855. PubMed PMID: 30478861; PMCID: PMC7379495.

20. Yanagisawa R, He C, Asai A, Hellwig M, Henle T, Toda M. The Impacts of Cholesterol, Oxysterols, and Cholesterol Lowering Dietary Compounds on the Immune System. Int J Mol Sci. 2022;23(20). Epub 20221013. doi: 10.3390/ijms232012236. PubMed PMID: 36293093; PMCID: PMC9602982.

21. Brown MS, Goldstein JL. The SREBP pathway: regulation of cholesterol metabolism by proteolysis of a membrane-bound transcription factor. Cell. 1997;89(3):331–40. doi: 10.1016/s0092-8674(00)80213-5. PubMed PMID: 9150132.

22. Repa JJ, Liang G, Ou J, Bashmakov Y, Lobaccaro JM, Shimomura I, Shan B, Brown MS, Goldstein JL, Mangelsdorf DJ. Regulation of mouse sterol regulatory element-binding protein-1c gene (SREBP-1c) by oxysterol receptors, LXRalpha and LXRbeta. Genes Dev. 2000;14(22):2819–30. doi: 10.1101/gad.844900. PubMed PMID: 11090130; PMCID: PMC317055.

23. Bjorkhem I. Do oxysterols control cholesterol homeostasis? J Clin Invest. 2002;110(6):725–30. doi: 10.1172/JCI16388. PubMed PMID: 12235099; PMCID: PMC151135.

24. Guillemot-Legris O, Mutemberezi V, Cani PD, Muccioli GG. Obesity is associated with changes in oxysterol metabolism and levels in mice liver, hypothalamus, adipose tissue and plasma. Scientific Reports. 2016;6. doi: 10.1038/srep19694. PubMed PMID: WOS:000368686700001.

25. Hintze KJ, Benninghoff AD, Ward RE. Formulation of the Total Western Diet (TWD) as a basal diet for rodent cancer studies. J Agric Food Chem. 2012;60(27):6736–42. Epub 20120124. doi: 10.1021/jf204509a. PubMed PMID: 22224871.

26. Monsanto SP, Hintze KJ, Ward RE, Larson DP, Lefevre M, Benninghoff AD. The new total Western diet for rodents does not induce an overweight phenotype or alter parameters of metabolic syndrome in mice. Nutr Res. 2016;36(9):1031–44. Epub 20160606. doi: 10.1016/j.nutres.2016.06.002. PubMed PMID: 27632924.

27. Xu Z, McClure ST, Appel LJ. Dietary Cholesterol Intake and Sources among U.S Adults: Results from National Health and Nutrition Examination Surveys (NHANES), 2001-2014. Nutrients. 2018;10(6). doi: 10.3390/nu10060771. PubMed PMID: WOS:000436507200115.

28. Maldonado-Pereira L, Barnaba C, Medina-Meza IG. Dietary exposure assessment of infant formula and baby foods’ oxidized lipids in the US population. Food Chem Toxicol. 2023;172:113552. Epub 20221209. doi: 10.1016/j.fct.2022.113552. PubMed PMID: 36502995.

29. Bushnell B. BBDuk Guide-DOE Joint Genome Institute. Computer software] https://jgidoegov/data-and-tools/bbtools/bb-tools-user-guide/bbduk-guide. 2021.

30. UniProt: the universal protein knowledgebase. Nucleic acids research. 2017;45(D1):D158–D69.

31. Franzosa EA, McIver LJ, Rahnavard G, Thompson LR, Schirmer M, Weingart G, Lipson KS, Knight R, Caporaso JG, Segata N, Huttenhower C. Species-level functional profiling of metagenomes and metatranscriptomes. Nat Methods. 2018;15(11):962–8. Epub 20181030. doi: 10.1038/s41592-018-0176-y. PubMed PMID: 30377376; PMCID: PMC6235447.

32. Caspi R, Billington R, Ferrer L, Foerster H, Fulcher CA, Keseler IM, Kothari A, Krummenacker M, Latendresse M, Mueller LA, Ong Q, Paley S, Subhraveti P, Weaver DS, Karp PD. The MetaCyc database of metabolic pathways and enzymes and the BioCyc collection of pathway/genome databases. Nucleic Acids Res. 2016;44(D1):D471–80. Epub 20151102. doi: 10.1093/nar/gkv1164. PubMed PMID: 26527732; PMCID: PMC4702838.

33. Blanco-Miguez A, Beghini F, Cumbo F, McIver LJ, Thompson KN, Zolfo M, Manghi P, Dubois L, Huang KD, Thomas AM, Nickols WA, Piccinno G, Piperni E, Puncochar M, Valles-Colomer M, Tett A, Giordano F, Davies R, Wolf J,…, Segata N. Extending and improving metagenomic taxonomic profiling with uncharacterized species using MetaPhlAn 4. Nat Biotechnol. 2023;41(11):1633–44. Epub 20230223. doi: 10.1038/s41587-023-01688-w. PubMed PMID: 36823356; PMCID: PMC10635831.

34. Wood DE, Lu J, Langmead B. Improved metagenomic analysis with Kraken 2. Genome Biol. 2019;20(1):257. Epub 20191128. doi: 10.1186/s13059-019-1891-0. PubMed PMID: 31779668; PMCID: PMC6883579.

35. Lu J, Breitwieser FP, Thielen P, Salzberg SL. Bracken: estimating species abundance in metagenomics data. PeerJ Comput Sci. 2017;3. Epub 20170102. doi: 10.7717/peerj-cs.104. PubMed PMID: 40271438; PMCID: PMC12016282.

36. Chen S, Zhou Y, Chen Y, Gu J. fastp: an ultra-fast all-in-one FASTQ preprocessor. Bioinformatics. 2018;34(17):i884–i90. doi: 10.1093/bioinformatics/bty560. PubMed PMID: 30423086; PMCID: PMC6129281.

37. Howe KL, Achuthan P, Allen J, Allen J, Alvarez-Jarreta J, Amode MR, Armean IM, Azov AG, Bennett R, Bhai J, Billis K, Boddu S, Charkhchi M, Cummins C, Da Rin Fioretto L, Davidson C, Dodiya K, El Houdaigui B, Fatima R,…, Flicek P. Ensembl 2021. Nucleic Acids Res. 2021;49(D1):D884–D91. doi: 10.1093/nar/gkaa942. PubMed PMID: 33137190; PMCID: PMC7778975.

38. Langmead B, Salzberg SL. Fast gapped-read alignment with Bowtie 2. Nat Methods. 2012;9(4):357–9. Epub 20120304. doi: 10.1038/nmeth.1923. PubMed PMID: 22388286; PMCID: PMC3322381.

39. Mallick H, Rahnavard A, McIver LJ, Ma S, Zhang Y, Nguyen LH, Tickle TL, Weingart G, Ren B, Schwager EH, Chatterjee S, Thompson KN, Wilkinson JE, Subramanian A, Lu Y, Waldron L, Paulson JN, Franzosa EA, Bravo HC, Huttenhower C. Multivariable association discovery in population-scale meta-omics studies. PLoS Comput Biol. 2021;17(11):e1009442. Epub 20211116. doi: 10.1371/journal.pcbi.1009442. PubMed PMID: 34784344; PMCID: PMC8714082.

40. Segata N, Izard J, Waldron L, Gevers D, Miropolsky L, Garrett WS, Huttenhower C. Metagenomic biomarker discovery and explanation. Genome Biol. 2011;12(6):R60. Epub 20110624. doi: 10.1186/gb-2011-12-6-r60. PubMed PMID: 21702898; PMCID: PMC3218848.

41. Di Tommaso P, Chatzou M, Floden EW, Barja PP, Palumbo E, Notredame C. Nextflow enables reproducible computational workflows. Nat Biotechnol. 2017;35(4):316–9. doi: 10.1038/nbt.3820. PubMed PMID: 28398311.

42. Carreon EJT. A Role for Adenomatous Polyposis Coli in Cellular Response to UV: University of Kansas; 2025.

43. Li B, Dewey CN. RSEM: accurate transcript quantification from RNA-Seq data with or without a reference genome. BMC Bioinformatics. 2011;12:323. Epub 20110804. doi: 10.1186/1471-2105-12-323. PubMed PMID: 21816040; PMCID: PMC3163565.

44. Team RC. RA language and environment for statistical computing, R Foundation for Statistical. Computing. 2020.

45. Ritchie ME, Phipson B, Wu D, Hu Y, Law CW, Shi W, Smyth GK. limma powers differential expression analyses for RNA-sequencing and microarray studies. Nucleic acids research. 2015;43(7):e47-e.

46. Stephens M. False discovery rates: a new deal. Biostatistics. 2017;18(2):275–94.

47. Wickham H. Elegant graphics for data analysis. Springer; 2016.

48. Wu T, Hu E, Xu S, Chen M, Guo P, Dai Z, Feng T, Zhou L, Tang W, Zhan L, Fu X, Liu S, Bo X, Yu G. clusterProfiler 4.0: A universal enrichment tool for interpreting omics data. Innovation (Camb). 2021;2(3):100141. Epub 20210701. doi: 10.1016/j.xinn.2021.100141. PubMed PMID: 34557778; PMCID: PMC8454663.

49. Kanehisa M, Goto S. KEGG: kyoto encyclopedia of genes and genomes. Nucleic Acids Res. 2000;28(1):27–30. doi: 10.1093/nar/28.1.27. PubMed PMID: 10592173; PMCID: PMC102409.

50. Gillespie M, Jassal B, Stephan R, Milacic M, Rothfels K, Senff-Ribeiro A, Griss J, Sevilla C, Matthews L, Gong C, Deng C, Varusai T, Ragueneau E, Haider Y, May B, Shamovsky V, Weiser J, Brunson T, Sanati N,…, D’Eustachio P. The reactome pathway knowledgebase 2022. Nucleic Acids Res. 2022;50(D1):D687-D92. doi: 10.1093/nar/gkab1028. PubMed PMID: 34788843; PMCID: PMC8689983.

51. Liberzon A, Birger C, Thorvaldsdottir H, Ghandi M, Mesirov JP, Tamayo P. The Molecular Signatures Database (MSigDB) hallmark gene set collection. Cell Syst. 2015;1(6):417–25. doi: 10.1016/j.cels.2015.12.004. PubMed PMID: 26771021; PMCID: PMC4707969.

52. Dolgalev I. msigdbr: MSigDB gene sets for multiple organisms in a tidy data format. R package version. 2022;7(1):532–2.

53. Korotkevich G, Sukhov V, Budin N, Shpak B, Artyomov MN, Sergushichev A. Fast gene set enrichment analysis. biorxiv. 2016:060012.

54. Badia IMP, Velez Santiago J, Braunger J, Geiss C, Dimitrov D, Muller-Dott S, Taus P, Dugourd A, Holland CH, Ramirez Flores RO, Saez-Rodriguez J. decoupleR: ensemble of computational methods to infer biological activities from omics data. Bioinform Adv. 2022;2(1):vbac016. Epub 20220308. doi: 10.1093/bioadv/vbac016. PubMed PMID: 36699385; PMCID: PMC9710656.

55. Garcia-Alonso L, Holland CH, Ibrahim MM, Turei D, Saez-Rodriguez J. Benchmark and integration of resources for the estimation of human transcription factor activities. Genome Res. 2019;29(8):1363–75. Epub 20190724. doi: 10.1101/gr.240663.118. PubMed PMID: 31340985; PMCID: PMC6673718.

56. Garcia-Alonso L, Holland CH, Ibrahim MM, Turei D, Saez-Rodriguez J. Corrigendum: Benchmark and integration of resources for the estimation of human transcription factor activities. Genome Res. 2021;31(4):745. doi: 10.1101/gr.275408.121. PubMed PMID: 33795376; PMCID: PMC8015848.

57. Hunter JD. Matplotlib: A 2D graphics environment. Computing in science & engineering. 2007;9(3):90–5.

58. Kolde R. Pheatmap: pretty heatmaps. R package version. 2019;1(2):726.

59. Stojanov S, Berlec A, Strukelj B. The Influence of Probiotics on the Firmicutes/Bacteroidetes Ratio in the Treatment of Obesity and Inflammatory Bowel disease. MICROORGANISMS. 2020;8. doi: 10.3390/microorganisms8111715. PubMed PMID: WOS:000593194600001.

60. Yang Z, Tang C, Sun X, Wu Z, Zhu X, Cui Q, Zhang R, Zhang X, Su Y, Mao Y. Protective effects of SCFAs on organ injury and gut microbiota modulation in heat-stressed rats. Annals of Microbiology. 2024;74(1):6.

61. Shetty SA, Zuffa S, Bui TPN, Aalvink S, Smidt H, De Vos WM. Reclassification of Eubacterium hallii as Anaerobutyricum hallii gen. nov., comb. nov., and description of Anaerobutyricum soehngenii sp. nov., a butyrate and propionate-producing bacterium from infant faeces. Int J Syst Evol Microbiol. 2018;68(12):3741–6. Epub 20181023. doi: 10.1099/ijsem.0.003041. PubMed PMID: 30351260.

62. Gilijamse PW, Hartstra AV, Levin E, Wortelboer K, Serlie MJ, Ackermans MT, Herrema H, Nederveen AJ, Imangaliyev S, Aalvink S, Sommer M, Levels H, Stroes ESG, Groen AK, Kemper M, de Vos WM, Nieuwdorp M, Prodan A. Treatment with Anaerobutyricum soehngenii: a pilot study of safety and dose-response effects on glucose metabolism in human subjects with metabolic syndrome. NPJ Biofilms Microbiomes. 2020;6(1):16. Epub 20200327. doi: 10.1038/s41522-020-0127-0. PubMed PMID: 32221294; PMCID: PMC7101376.

63. Hosomi K, Saito M, Park J, Murakami H, Shibata N, Ando M, Nagatake T, Konishi K, Ohno H, Tanisawa K, Mohsen A, Chen YA, Kawashima H, Natsume-Kitatani Y, Oka Y, Shimizu H, Furuta M, Tojima Y, Sawane K, Saika A, Kondo S, Yonejima Y, Takeyama H, Matsutani A, Mizuguchi K, Miyachi M, Kunisawa J. Oral administration of Blautia wexlerae ameliorates obesity and type 2 diabetes via metabolic remodeling of the gut microbiota. Nat Commun. 2022;13(1):4477. Epub 20220818. doi: 10.1038/s41467-022-32015-7. PubMed PMID: 35982037; PMCID: PMC9388534.

64. Parker BJ, Wearsch PA, Veloo ACM, Rodriguez-Palacios A. The Genus Alistipes: Gut Bacteria With Emerging Implications to Inflammation, Cancer, and Mental Health. Front Immunol. 2020;11:906. Epub 20200609. doi: 10.3389/fimmu.2020.00906. PubMed PMID: 32582143; PMCID: PMC7296073.

65. Wu TR, Lin CS, Chang CJ, Lin TL, Martel J, Ko YF, Ojcius DM, Lu CC, Young JD, Lai HC. Gut commensal Parabacteroides goldsteinii plays a predominant role in the anti-obesity effects of polysaccharides isolated from Hirsutella sinensis. Gut. 2019;68(2):248–62. Epub 20180714. doi: 10.1136/gutjnl-2017-315458. PubMed PMID: 30007918.

66. Lee AH, Rodriguez Jimenez DM, Meisel M. Limosilactobacillus reuteri - a probiotic gut commensal with contextual impact on immunity. Gut Microbes. 2025;17(1):2451088. Epub 20250117. doi: 10.1080/19490976.2025.2451088. PubMed PMID: 39825615; PMCID: PMC12716054.

67. Lu J, Rincon N, Wood DE, Breitwieser FP, Pockrandt C, Langmead B, Salzberg SL, Steinegger M. Metagenome analysis using the Kraken software suite. Nat Protoc. 2022;17(12):2815–39. Epub 20220928. doi: 10.1038/s41596-022-00738-y. PubMed PMID: 36171387; PMCID: PMC9725748.

68. Lu J, Rincon N, Wood DE, Breitwieser FP, Pockrandt C, Langmead B, Salzberg SL, Steinegger M. Author Correction: Metagenome analysis using the Kraken software suite. Nat Protoc. 2026;21(2):872. doi: 10.1038/s41596-024-01064-1. PubMed PMID: 39210095.

69. Drula E, Garron ML, Dogan S, Lombard V, Henrissat B, Terrapon N. The carbohydrate-active enzyme database: functions and literature. Nucleic Acids Res. 2022;50(D1):D571–D7. doi: 10.1093/nar/gkab1045. PubMed PMID: 34850161; PMCID: PMC8728194.

70. Irving SE, Choudhury NR, Corrigan RM. The stringent response and physiological roles of (pp)pGpp in bacteria. Nat Rev Microbiol. 2021;19(4):256–71. Epub 20201104. doi: 10.1038/s41579-020-00470-y. PubMed PMID: 33149273.

71. Heuston S, Begley M, Gahan CGM, Hill C. Isoprenoid biosynthesis in bacterial pathogens. Microbiology (Reading). 2012;158(Pt 6):1389–401. Epub 20120330. doi: 10.1099/mic.0.051599-0. PubMed PMID: 22466083.

72. Flamholz A, Noor E, Bar-Even A, Liebermeister W, Milo R. Glycolytic strategy as a tradeoff between energy yield and protein cost. Proc Natl Acad Sci U S A. 2013;110(24):10039–44. Epub 20130429. doi: 10.1073/pnas.1215283110. PubMed PMID: 23630264; PMCID: PMC3683749.

73. Heumuller-Klug S, Sticht C, Kaiser K, Wink E, Hagl C, Wessel L, Schafer KH. Degradation of intestinal mRNA: a matter of treatment. World J Gastroenterol. 2015;21(12):3499–508. doi: 10.3748/wjg.v21.i12.3499. PubMed PMID: 25834314; PMCID: PMC4375571.

74. Kruit JK, Groen AK, van Berkel TJ, Kuipers F. Emerging roles of the intestine in control of cholesterol metabolism. World J Gastroenterol. 2006;12(40):6429–39. doi: 10.3748/wjg.v12.i40.6429. PubMed PMID: 17072974; PMCID: PMC4100631.

75. Radhakrishnan A, Ikeda Y, Kwon HJ, Brown MS, Goldstein JL. Sterol-regulated transport of SREBPs from endoplasmic reticulum to Golgi: oxysterols block transport by binding to Insig. Proc Natl Acad Sci U S A. 2007;104(16):6511–8. Epub 20070411. doi: 10.1073/pnas.0700899104. PubMed PMID: 17428920; PMCID: PMC1851665.

76. He Y, Dong Y, Zhang X, Ding Z, Song Y, Huang X, Chen S, Wang Z, Ni Y, Ding L. Lipid Droplet-Related PLIN2 in CD68+ Tumor-Associated Macrophage of Oral Squamous Cell Carcinoma: Implications for Cancer Prognosis and Immunotherapy. FRONTIERS IN ONCOLOGY. 2022;12. doi: 10.3389/fonc.2022.824235. PubMed PMID: WOS:000778651700001.

77. Fink K, Grandvaux N. STAT2 and IRF9: Beyond ISGF3. JAKSTAT. 2013;2(4):e27521. Epub 20131218. doi: 10.4161/jkst.27521. PubMed PMID: 24498542; PMCID: PMC3906322.

78. Platanias LC. Mechanisms of type-I- and type-II-interferon-mediated signalling. Nat Rev Immunol. 2005;5(5):375–86. doi: 10.1038/nri1604. PubMed PMID: 15864272.

79. Rauch I, Rosebrock F, Hainzl E, Heider S, Majoros A, Wienerroither S, Strobl B, Stockinger S, Kenner L, Muller M, Decker T. Noncanonical Effects of IRF9 in Intestinal Inflammation: More than Type I and Type III Interferons. Mol Cell Biol. 2015;35(13):2332–43. Epub 20150427. doi: 10.1128/MCB.01498-14. PubMed PMID: 25918247; PMCID: PMC4456449.

80. Juul F, Martinez-Steele E, Parekh N, Monteiro CA, Chang VW. Ultra-processed food consumption and excess weight among US adults. British Journal of Nutrition. 2018;120(1):90–100. doi: 10.1017/s0007114518001046. PubMed PMID: WOS:000436257400012.

81. Ntambi JM, Miyazaki M. Regulation of stearoyl-CoA desaturases and role in metabolism. Prog Lipid Res. 2004;43(2):91–104. doi: 10.1016/s0163-7827(03)00039-0. PubMed PMID: 14654089.

82. Russell DW. The enzymes, regulation, and genetics of bile acid synthesis. Annu Rev Biochem. 2003;72:137–74. Epub 20030116. doi: 10.1146/annurev.biochem.72.121801.161712. PubMed PMID: 12543708.

83. Wahlstrom A, Sayin SI, Marschall HU, Backhed F. Intestinal Crosstalk between Bile Acids and Microbiota and Its Impact on Host Metabolism. Cell Metab. 2016;24(1):41–50. Epub 20160616. doi: 10.1016/j.cmet.2016.05.005. PubMed PMID: 27320064.

84. Devlin AS, Fischbach MA. A biosynthetic pathway for a prominent class of microbiota-derived bile acids. Nat Chem Biol. 2015;11(9):685–90. Epub 20150720. doi: 10.1038/nchembio.1864. PubMed PMID: 26192599; PMCID: PMC4543561.

85. Cuzzocrea S, Ianaro A, Wayman NS, Mazzon E, Pisano B, Dugo L, Serraino I, Di Paola R, Chatterjee PK, Di Rosa M, Caputi AP, Thiemermann C. The cyclopentenone prostaglandin 15-deoxy-delta(12,14)-PGJ2 attenuates the development of colon injury caused by dinitrobenzene sulphonic acid in the rat. Br J Pharmacol. 2003;138(4):678–88. doi: 10.1038/sj.bjp.0705077. PubMed PMID: 12598422; PMCID: PMC1573694.

86. Straus DS, Pascual G, Li M, Welch JS, Ricote M, Hsiang C-H, Sengchanthalangsy LL, Ghosh G, Glass CK. 15-Deoxy-Δ12, 14-prostaglandin J2 inhibits multiple steps in the NF-κB signaling pathway. Proceedings of the National Academy of Sciences. 2000;97(9):4844–9.

87. Wlodarska M, Luo C, Kolde R, d’Hennezel E, Annand JW, Heim CE, Krastel P, Schmitt EK, Omar AS, Creasey EA, Garner AL, Mohammadi S, O’Connell DJ, Abubucker S, Arthur TD, Franzosa EA, Huttenhower C, Murphy LO, Haiser HJ,…, Xavier RJ. Indoleacrylic Acid Produced by Commensal Peptostreptococcus Species Suppresses Inflammation. Cell Host Microbe. 2017;22(1):25–37 e6. doi: 10.1016/j.chom.2017.06.007. PubMed PMID: 28704649; PMCID: PMC5672633.

88. Cervantes-Barragan L, Chai JN, Tianero MD, Di Luccia B, Ahern PP, Merriman J, Cortez VS, Caparon MG, Donia MS, Gilfillan S, Cella M, Gordon JI, Hsieh CS, Colonna M. Lactobacillus reuteri induces gut intraepithelial CD4(+)CD8alphaalpha(+) T cells. Science. 2017;357(6353):806–10. Epub 20170803. doi: 10.1126/science.aah5825. PubMed PMID: 28775213; PMCID: PMC5687812.

89. Pegg AE. Mammalian polyamine metabolism and function. IUBMB Life. 2009;61(9):880-94. doi: 10.1002/iub.230. PubMed PMID: 19603518; PMCID: PMC2753421.

90. Lorbek G, Perse M, Horvat S, Bjorkhem I, Rozman D. Sex differences in the hepatic cholesterol sensing mechanisms in mice. Molecules. 2013;18(9):11067–85. Epub 20130910. doi: 10.3390/molecules180911067. PubMed PMID: 24025456; PMCID: PMC6270450.

91. Schultz JR, Tu H, Luk A, Repa JJ, Medina JC, Li L, Schwendner S, Wang S, Thoolen M, Mangelsdorf DJ, Lustig KD, Shan B. Role of LXRs in control of lipogenesis. Genes Dev. 2000;14(22):2831–8. doi: 10.1101/gad.850400. PubMed PMID: 11090131; PMCID: PMC317060.

92. Grefhorst A, Elzinga BM, Voshol PJ, Plosch T, Kok T, Bloks VW, van der Sluijs FH, Havekes LM, Romijn JA, Verkade HJ, Kuipers F. Stimulation of lipogenesis by pharmacological activation of the liver X receptor leads to production of large, triglyceride-rich very low density lipoprotein particles. J Biol Chem. 2002;277(37):34182–90. Epub 20020703. doi: 10.1074/jbc.M204887200. PubMed PMID: 12097330.

93. Kilvington A, Barnaba C, Rajasekaran S, Laurens Leimanis ML, Medina-Meza IG. Lipid profiling and dietary assessment of infant formulas reveal high intakes of major cholesterol oxidative product (7-ketocholesterol). Food Chem. 2021;354:129529. Epub 20210309. doi: 10.1016/j.foodchem.2021.129529. PubMed PMID: 33761334.

94. DeBose-Boyd RA. Feedback regulation of cholesterol synthesis: sterol-accelerated ubiquitination and degradation of HMG CoA reductase. Cell Res. 2008;18(6):609–21. doi: 10.1038/cr.2008.61. PubMed PMID: 18504457; PMCID: PMC2742364.

95. Sever N, Yang T, Brown MS, Goldstein JL, DeBose-Boyd RA. Accelerated degradation of HMG CoA reductase mediated by binding of insig-1 to its sterol-sensing domain. Mol Cell. 2003;11(1):25–33. doi: 10.1016/s1097-2765(02)00822-5. PubMed PMID: 12535518.

96. Wang F, Kohan AB, Lo CM, Liu M, Howles P, Tso P. Apolipoprotein A-IV: a protein intimately involved in metabolism. J Lipid Res. 2015;56(8):1403–18. Epub 20150201. doi: 10.1194/jlr.R052753. PubMed PMID: 25640749; PMCID: PMC4513983.

97. Montgomery TL, Eckstrom K, Lile KH, Caldwell S, Heney ER, Lahue KG, D’Alessandro A, Wargo MJ, Krementsov DN. Lactobacillus reuteri tryptophan metabolism promotes host susceptibility to CNS autoimmunity. Microbiome. 2022;10(1):198. Epub 20221123. doi: 10.1186/s40168-022-01408-7. PubMed PMID: 36419205; PMCID: PMC9685921.

98. Scott SA, Fu J, Chang PV. Microbial tryptophan metabolites regulate gut barrier function via the aryl hydrocarbon receptor. Proc Natl Acad Sci U S A. 2020;117(32):19376–87. Epub 20200727. doi: 10.1073/pnas.2000047117. PubMed PMID: 32719140; PMCID: PMC7431026.

99. Nakamura A, Kurihara S, Takahashi D, Ohashi W, Nakamura Y, Kimura S, Onuki M, Kume A, Sasazawa Y, Furusawa Y, Obata Y, Fukuda S, Saiki S, Matsumoto M, Hase K. Symbiotic polyamine metabolism regulates epithelial proliferation and macrophage differentiation in the colon. Nat Commun. 2021;12(1):2105. Epub 20210408. doi: 10.1038/s41467-021-22212-1. PubMed PMID: 33833232; PMCID: PMC8032791.

100. Rao JN, Xiao L, Wang JY. Polyamines in Gut Epithelial Renewal and Barrier Function. Physiology (Bethesda). 2020;35(5):328–37. doi: 10.1152/physiol.00011.2020. PubMed PMID: 32783609; PMCID: PMC7642847.

